# A Benchmark of Modern Statistical Phasing Methods

**DOI:** 10.1101/2025.06.24.660794

**Authors:** Andrew T. Beck, Hyun Min Kang, Sebastian Zöllner

## Abstract

Modern statistical phasing methods efficiently and accurately infer haplotype phase in large genomic samples, and their performance is critical for downstream analyses. However, rigorously evaluating phasing accuracy remains challenging. Here we demonstrate the use of synthetic diploids generated from male X chromosome sequences to benchmark and compare three commonly-used phasing methods: Beagle 5.4, SHAPEIT 4, and Eagle v2.4.1. Our evaluation reveals highly correlated error rates across all methods, although we observe distinct variation among methods within error types. Specifically, Eagle v2.4.1 is characterized by a higher frequency of switch errors, whereas SHAPEIT 4 displays a higher rate of flip (double-switch) errors. These patterns are consistent across diverse populations. Additionally, we observe an enrichment of errors at both CpG sites and rare variant sites, particularly for flip errors. Validating these results in autosomal trio data, we observe similar trends between the methods and error patterns, though with an increased flip error rate, a discrepancy that may be biased by genotype error.

## 1 Introduction

In tandem with the explosive growth of genome sequencing sample sizes, modern population-based statistical phasing methods have evolved to handle the complex task of efficiently constructing phased haplotypes for many samples [1–3]. Phased haplotypes are essential for a variety of downstream applications, ranging from imputation, which increases statistical power in association studies by improving SNP coverage [4], to the identification of compound heterozygous events [5, 6]. Modern statistical phasing methods such as Eagle [1], Beagle [2], and SHAPEIT [3] are able to efficiently determine the haplotype phase in thousands to millions of samples with high precision, and the increasing availability of larger and more diverse reference panels has also led to improvements in phasing accuracy [7]. Despite these improvements in both the phasing methods and the sample sizes, statistical phasing remains a challenging task, and modern phasing algorithms still introduce thousands of errors in each phased genome [8]. These errors impact downstream analyses, and therefore a comprehensive analysis of their frequency and the genomic contexts in which they occur more frequently is of considerable interest.

Previous efforts to benchmark phasing methods have relied on either genotyped trios, or genomes phased using a combination of multiple sequencing platforms and experimental techniques [4, 8]. While pedigree information allows for the construction of phased haplotypes under the principles of Mendelian inheritance, switch and flip error rates estimated from trio data are inflated due to genotyping error [9]. In addition, the cost of obtaining parent-child samples often precludes their widespread use in genetic studies, whereas large samples of unrelated individuals are readily available [10–12]. While these vast datasets of unrelated individuals offer the scale needed to characterize phasing errors in granular detail, the lack of gold-standard ground truth haplotypes prevents a direct evaluation of performance. Thus, previous efforts to benchmark phasing methods using these large samples of unrelated individuals have relied on imputation quality as a downstream metric of phasing quality [4, 13]. Imputation accuracy is a reflection of the underlying phasing quality and thus allows for a comparison across methods, but it does not allow for a detailed evaluation of the genomic context in which phasing errors occur as the number and location of phasing errors are unknown.

To directly evaluate statistical phasing methods on unrelated individuals and identify the quantity and location of phasing errors, known haplotypes are required. While haplotype phase is initially unknown for most sequenced chromosomes, the male X chromosome is hemizygous outside of the pseudo-autosomal region (PAR), and thus the phase is readily available from the genotypes. This known phase allows us to construct synthetic diploids by sampling pairs of male X chromosomes [14]. These synthetic diploids can then be phased and the results can then be compared to the known haplotypes, enabling both the estimation of phasing error rates and a characterization of the locations where these errors occur. Large studies of unrelated individuals across diverse populations are readily available, and thus synthetic diploids offer an avenue by which the rate of statistical phasing errors can be evaluated and characterized with regards to the genomic contexts in which they occur.

To illustrate the utility of this approach, we benchmark three modern statistical phasing software implementations (Beagle 5.4, Eagle 2.4.1, and SHAPEIT 4), constructing synthetic diploids from male X chromosomes sampled from 2,504 unrelated individuals in the 1000 Genomes Project (1kGP) deep sequence release [12]. Each synthetic diploid was phased individually with all three algorithms and the inferred haplotypes were then compared to the original haplotypes to identify phasing errors. To better understand the drivers of such errors, it is useful to distinguish between switch errors (haplotype changes incorrectly from matching one paternal haplotype to matching the other) and flip errors (a single variant’s phase is incorrect relative to its neighbors) [9]. Our evaluation reveals that while the distributions of errors are highly correlated among the three methods, they exhibit distinct error profiles: Eagle v2.4.1 displays a higher frequency of switch errors, whereas SHAPEIT 4 introduces flip errors at a higher rate. When examining genomic drivers, we found that flip errors were primarily enriched at CpG sites and rare variants, while switch errors were enriched in regions of high recombination. To evaluate the generalizability of these X-chromosome benchmarks to autosomal data, we phased 602 Mendelian-resolved trio probands. We observe that while switch error rates are representative across both autosomal and X genomic contexts, flip error rates are elevated in the autosomes, a discrepancy potentially driven by genotype error [9].

## 2 Results

We simulated synthetic diploids with known phase by sampling male X chromosomes from the 1kGP dataset and phased each synthetic diploid sample using Eagle v2.4.1, Beagle 5.5 and SHAPEIT4 (as implemented in the SHAPEIT v5.0.1 software). We compared each method’s inferred chromosomes to the known chromosomes to identify switch and flip errors. To assess consistency across chromosomes, we rephased the autosomes of probands from each of the 602 trios of the extended 1kGP sample while ignoring the available pedigree information, comparing the results to pedigree-adjusted phased chromosomes.

### 2.1 Distributions of Errors in Synthetic Diploids

Across a mean of 68,930 heterozygous sites per sample, all algorithms showed a high degree of accuracy. The mean total error counts per sample were lowest for Beagle (197.7) and highest for Eagle (236.5), with SHAPEIT averaging 201.2 errors per sample. To account for individual variation in heterozygosity, we calculated a per-sample error rate for each synthetic diploid; the resulting average error rates across the samples were 0.0031 for both Beagle and SHAPEIT, and 0.0037 for Eagle. Despite a relatively small difference in the mean error rates, one-way repeated measures ANOVA indicated that the error rates differed significantly across the methods (*p <* 2.2 *×* 10*^−^*^16^), and all pairs of methods were found to be statistically significantly different.

Across individual samples, we observe substantial variation of the error rate within each method ranging from 4.05 *×* 10*^−^*^4^ to 5.13 *×* 10*^−^*^3^ errors per heterozygous position, with samples clustering by 1kGP population (Fig. A1). Comparing across populations, both the mean number of heterozygous sites per sample and error counts differed greatly, with AFR samples having on average the largest number of heterozygous positions (101673.52 heterozygous sites per sample), whereas EAS and SAS samples had an overall higher mean error count and mean error rate (Table 1). Comparing methods within populations we observed that in all populations Beagle had the lowest error rate, followed by SHAPEIT while Eagle had the largest error rate.

**Table 1:**
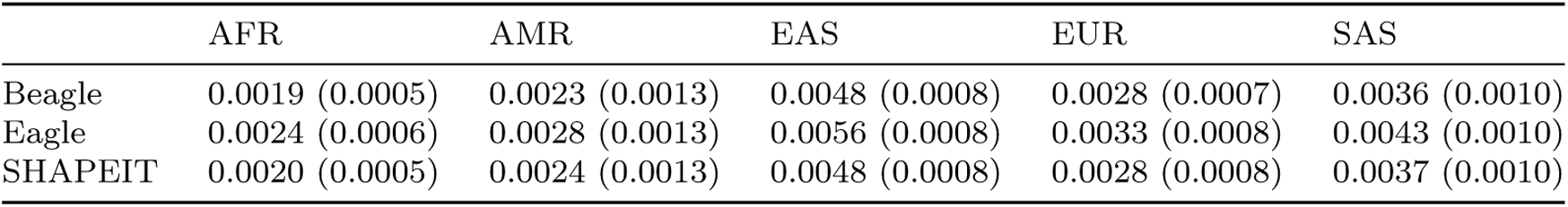
Phasing error rates stratified by 1kGP population. Mean (SD) errors per heterozygous site calculated across 200 synthetic diploids for each of the five 1000 Genomes Project populations.

|  | AFR | AMR | EAS | EUR | SAS |
| --- | --- | --- | --- | --- | --- |
| Beagle | 0.0019 (0.0005) | 0.0023 (0.0013) | 0.0048 (0.0008) | 0.0028 (0.0007) | 0.0036 (0.0010) |
| Eagle | 0.0024 (0.0006) | 0.0028 (0.0013) | 0.0056 (0.0008) | 0.0033 (0.0008) | 0.0043 (0.0010) |
| SHAPEIT | 0.0020 (0.0005) | 0.0024 (0.0013) | 0.0048 (0.0008) | 0.0028 (0.0008) | 0.0037 (0.0010) |

We next considered switch errors and flip errors separately. Overall, switch errors constituted 54.3% of all observed errors across all samples and methods, though this composition varied significantly by algorithm (*p <* 2.2 *×* 10*^−^*^16^). Specifically, switch errors accounted for 53% of Beagle’s total errors, 61% of Eagle’s, and 47% of SHAPEIT’s. When comparing average per-sample error rates, SHAPEIT introduced the fewest switch errors (0.0016 switches per heterozygous site), followed by Beagle (0.0017) and Eagle (0.0023). These rates correspond to an average of 98.3, 107.4, and 146.3 switches per sample, respectively. Similarly, flip rates showed a significant difference across the methods (*p <* 2.2 *×* 10*^−^*^16^). In contrast to switch errors, SHAPEIT yielded the highest flip rate (0.0016 flips per heterozygous site), while Beagle (0.0014) and Eagle (0.0014) demonstrated lower rates. These rates translate to an average of 102.9, 90.4, and 90.2 flips per sample for SHAPEIT, Beagle, and Eagle, respectively.

**Fig. 1:**
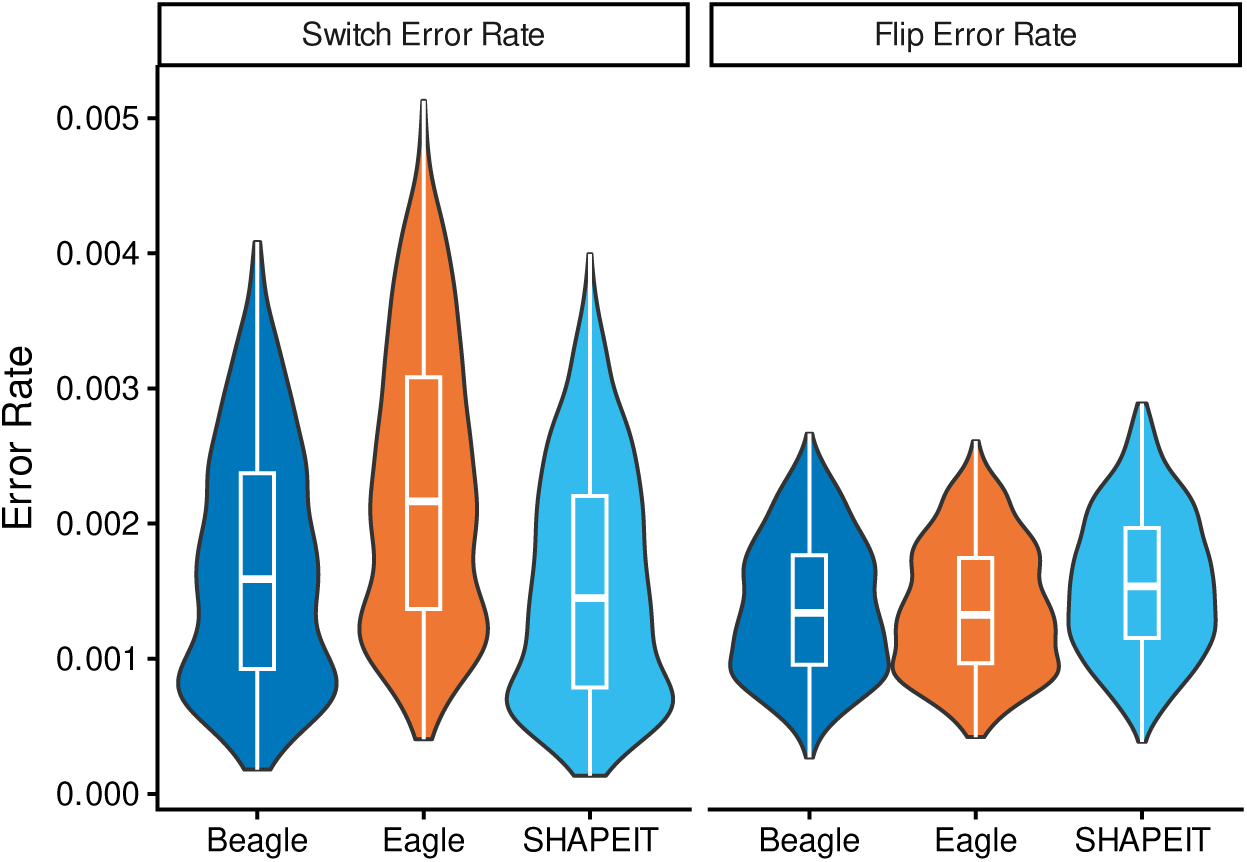
Switch and flip error rate distributions across 1000 synthetic X-chromosome diploids. Switch error rates varied significantly across all methods (*p <* 10*^−^*^15^), with Eagle exhibiting the highest mean rate. Conversely, SHAPEIT showed a significantly higher flip error rate than both Beagle and Eagle (*p <* 10*^−^*^15^), while flip error rates between Beagle and Eagle did not differ significantly (*p* = 0.42).

While all pairs of methods were found to have a significantly different switch rate (*p <* 0.0001), the difference between flip rates for Beagle and Eagle was not found to be significantly different (*p* = 0.72). This ranking of methods by both switch rate and flip rate was observed to be consistent across the five 1kGP populations. Switch rates and flip rates differed significantly among populations (*p <* 2.2 *×* 10*^−^*^16^ for both). For both error types, the EAS population exhibited the highest average rate, while the AFR population showed the lowest average rate. (Table 2).

**Table 2:** Distribution of switch and flip phasing errors by 1kGP population. Comparison of single switch error rates and flip (double switch) error rates across five 1kGP populations. Error rates are normalized per heterozygous position and averaged over 200 synthetic diploids.

|  | Switch Error Rates |  |  | Flip Error Rates |  |  |
| --- | --- | --- | --- | --- | --- | --- |
|  | Beagle | Eagle | SHAPEIT | Beagle | Eagle | SHAPEIT |
| AFR | 0.0009 (0.0003) | 0.0014 (0.0004) | 0.0008 (0.0003) | 0.0010 (0.0003) | 0.0010 (0.0003) | 0.0012 (0.0003) |
| AMR | 0.0012 (0.0008) | 0.0017 (0.0009) | 0.0011 (0.0008) | 0.0011 (0.0005) | 0.0011 (0.0004) | 0.0013 (0.0005) |
| EAS | 0.0028 (0.0005) | 0.0037 (0.0006) | 0.0027 (0.0005) | 0.0020 (0.0003) | 0.0019 (0.0003) | 0.0021 (0.0004) |
| EUR | 0.0015 (0.0004) | 0.0021 (0.0005) | 0.0014 (0.0004) | 0.0012 (0.0003) | 0.0012 (0.0003) | 0.0014 (0.0004) |
| SAS | 0.0020 (0.0006) | 0.0027 (0.0007) | 0.0019 (0.0006) | 0.0016 (0.0004) | 0.0016 (0.0004) | 0.0018 (0.0004) |

The number of total errors for each synthetic diploid was highly correlated between methods, with correlation coefficients ranging from 0.979 to 0.982. The number of switches and the number of flips were also highly correlated. Specific phasing errors were often shared between methods (i.e. they were the same type of errors occurring at the same location in at least two methods for the same sample). Across the 1000 synthetic diploids, 59.5% of the Beagle switches were shared with at least one other method, while 63.9% of the SHAPEIT switches were shared with at least one other method, and 39.6% of Eagle switches were shared with at least one other method. Overlap in flip errors were generally higher, where we observe that 65.8% of Beagle’s flips are shared with at least one other method, 57.1% of Eagle’s flips are shared with at least one other method, and 61% of SHAPEIT’s flips are shared with at least one other method.

The rate of errors for all three methods were heterogeneous across the X chromosome (Figure A2). Calculated in 1mb bins, average errors rates for switches range from 8 *·* 10*^−^*^5^ to 7 *·* 10*^−^*^3^ switches per heterozygous position while error rates for flips range from 6 *·* 10*^−^*^4^ to 7 *·* 10*^−^*^3^ flips per heterozygous position. Between methods, the distribution of both switch and flip rates across bins was highly correlated (*ρ ≈* 1). Nonetheless, Eagle’s average error rate was higher than SHAPEIT’s in all bins, and higher than Beagle’s in all but one bin. On average, the switch error rate observed for Eagle within a bin was 1.75 times that of Beagle’s and 2.18 times that observed for SHAPEIT (Supplementary Figure A3). In contrast, we observed a higher flip rate in most bins for SHAPEIT compared to Beagle and Eagle, with SHAPEIT’s flip rate in each bin being on average 1.14 times that of Beagle and 1.15 times that of Eagle (Supplementary Figure A4).

### 2.2 Genomic predictors of phasing error

To determine if context-dependent mutation rates influence phasing accuracy, we analyzed the distribution of switch and flip errors at CpG sites, which exhibit roughly 10-fold higher mutability than non-CpG contexts[15–19]. While CpG sites constituted only 14.2% of all heterozygous sites across the synthetic diploids, they accounted for a disproportionate share of phasing errors, representing approximately 24% of the total error burden across all three methods. This enrichment was notably more pronounced for flip errors than for switch errors. Approximately one-third of all flip errors occurred at CpG sites (Beagle: 34%, *OR* = 3.09; SHAPEIT: 34%, *OR* = 3.07; Eagle: 30%, *OR* = 2.60), representing a high degree of localized sensitivity. In contrast, switch error frequencies were only marginally elevated (Beagle: 16%, *OR* = 1.16; SHAPEIT: 15%, *OR* = 1.11; Eagle: 19%, *OR* = 1.43). These associations for both flips and switches were statistically significant across all three methods (*p <* 2.8 *×* 10*^−^*^33^). The patterns of enrichment were also found to be consistent across all 1kGP populations (Supplementary Figure A5).

We then examined the contribution of rare variants to phasing errors by comparing the distributions of minor allele frequencies (maf) at phasing errors to the maf distributions at heterozygous positions within each synthetic diploid. We categorized variants as either rare (maf *≤* 0.05), uncommon (0.05 *<* maf *≤* 0.1), or common (maf *>* 0.1). Across the synthetic diploids, rare, uncommon and common sites constituted an average of 6.02%, 8.53% and 85.5% of all heterozygous sites respectively. Among all errors, 39.3% occurred at rare sites (Supplementary Tables B2 and B3). The common odds ratio of the odds of a switch error occurring at a rare variant over the odds of it occurring at a non-rare site were 4.44, 7.25, and 4.15 for Beagle, Eagle and SHAPEIT, respectively. We observed a stronger enrichment of flips at rare variants, with odd ratios of flips occurring at rare sites compared to other sites of 20.02, 19.07, and 19.03 for Beagle, Eagle, and SHAPEIT, respectively. All rare variant enrichments for both error types were highly significant (*p <* 1 *×* 10*^−^*^300^).

To examine the relationship between recombination rate and phasing accuracy, we divided the X chromosome into non-overlapping 1 MB bins. Across the 755 resulting bins, we observed a positive correlation between average recombination rate and switch error rates (*r* = 0.78(*p <* 0.001), *r* = 0.45(*p <* 0.001), and *r* = 0.81(*p <* 0.001) for Beagle, Eagle, and SHAPEIT, respectively). In contrast, the correlation between recombination and flip error rates was negative, but not statistically significantly different from 0 (*r* = *−*0.11(*p* = 0.19), *r* = *−*0.06(*p* = 0.44), and *r* = *−*0.136(*p* = 0.17))

(Supplementary Figures A10, A11). To examine whether switches and flips were enriched in regions with the highest recombination rates, we assigned each error to a recombination rate decile (See methods). For Beagle and SHAPEIT, we observed that on average over 40% of switch errors occurred at heterozygous positions with recombination rates in the top decile, and over 30% of Eagle switch errors on average were also in this top decile. For flips we observed lower enrichment of errors in the top recombination rate decile, but in all methods on average over 15% of flip errors occurred at positions with recombination rates in the top decile (Supplementary Figures A12, A13).

### 2.3 Error Distributions in the Autosomes

To evaluate the robustness of our observations and comparisons of methods in chromosome X to other chromosomes, we rephased the autosomes of 602 trio probands from the 1kGP deep-sequence release while excluding their parents’ haplotypes. Across the autosomes, Beagle’s error rates varied from 0.0065 to 0.0105 errors per heterozygous site, Eagle’s from 0.0063 to 0.0105, and SHAPEIT’s from 0.0069 to 0.0109. These error rates were all higher than what we had observed in the X chromosome for the synthetic diploids (Beagle: 0.0031 errors per heterozygous site; Eagle: 0.0037; SHAPEIT: 0.0031). This difference was largely due to an increased flip rate in the autosomes relative to X, with switch rates in the autosomes aligning with what was observed in the X chromosome (Figure 3). Average flip rates in the autosomes varied between 0.0050 and 0.0067 flips per heterozygous site in Beagle, 0.0044 and 0.0060 in Eagle, and 0.0054 and 0.0072 in SHAPEIT, whereas the average flip rates in X were 0.0014, 0.0014, and 0.0016 for Beagle, Eagle, and SHAPEIT. In contrast, switch rates in the autosomes ranged from 0.0015 to 0.0038 switches per heterozygous site in Beagle, 0.0020 to 0.0045 in Eagle, and from 0.0015 to 0.0037 in SHAPEIT, with the corresponding average switch rates in X being 0.0017, 0.0023, and 0.0016 in Beagle, Eagle, and SHAPEIT. Despite differences in error rates observed across chromosomes, the rankings of methods was found to be consistent both across all autosomes and between proband and synthetic diploid results. In particular, in all chromosomes Eagle has a higher average switch error rate, while SHAPEIT and Beagle have similar average switch error rates. In contrast, SHAPEIT has a higher average flip rate in all chromosomes and Eagle has the lowest flip error rate.

**Fig. 2:**
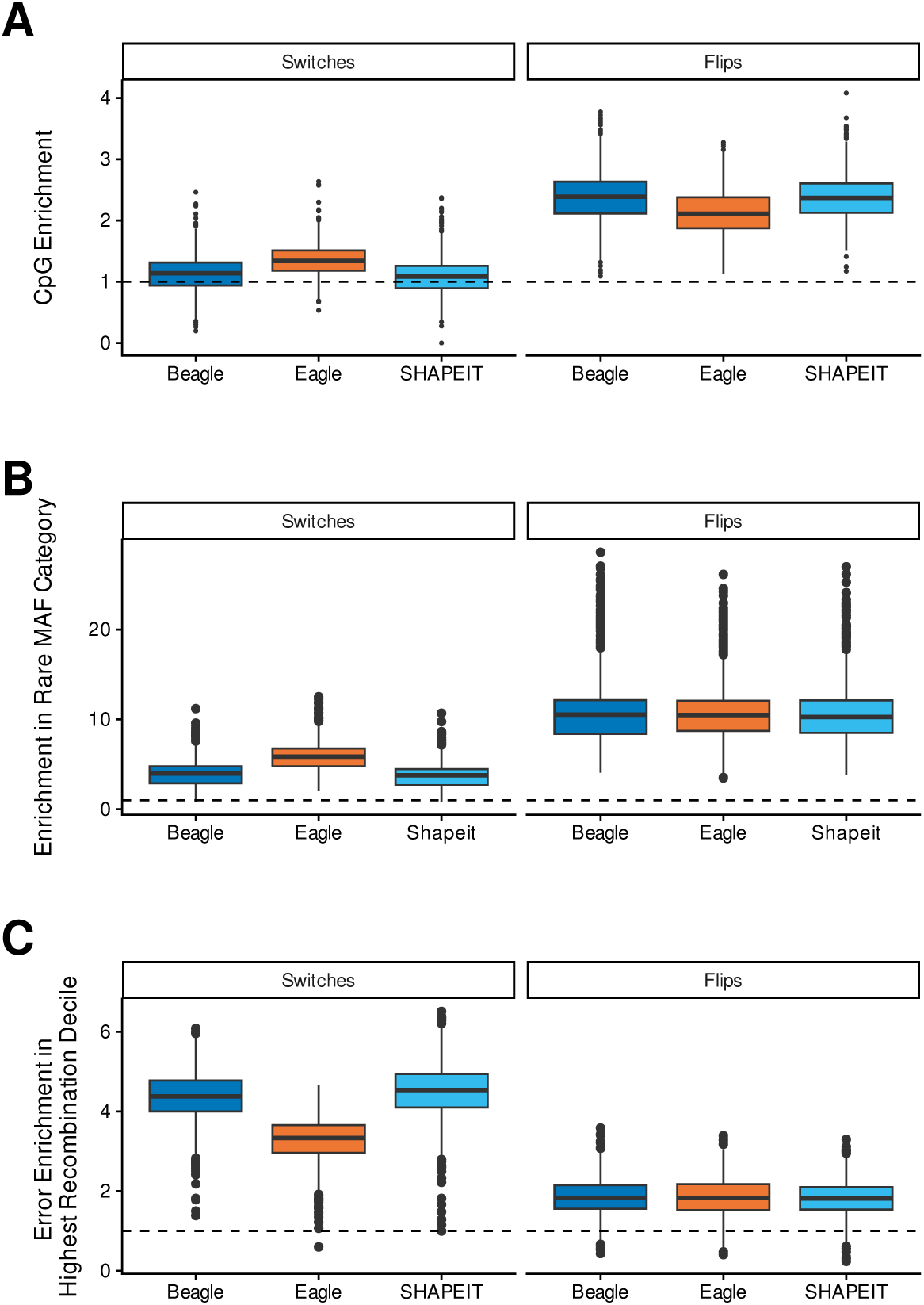
Box plots illustrating the distribution of switch and flip error enrichment across all individuals for three genomic features. For each feature, enrichment is defined as the proportion of errors occurring at the feature divided by the proportion of the individual’s total heterozygous sites located at the feature. **(A) CpG Sites:** Enrichment of errors occurring at CpG sites relative to the background proportion of heterozygous sites at CpGs. **(B) Rare Variants:** Enrichment of errors at rare variant sites (*MAF <* 0.05). **(C) High Recombination Rate:** Enrichment of errors at the highest decile of recombination rates. For each individual, heterozygous sites were mapped to the nearest recombination rate value; the enrichment represents the proportion of errors in the top 10th percentile of these values divided by the expected proportion (0.1).

**Fig. 3:**
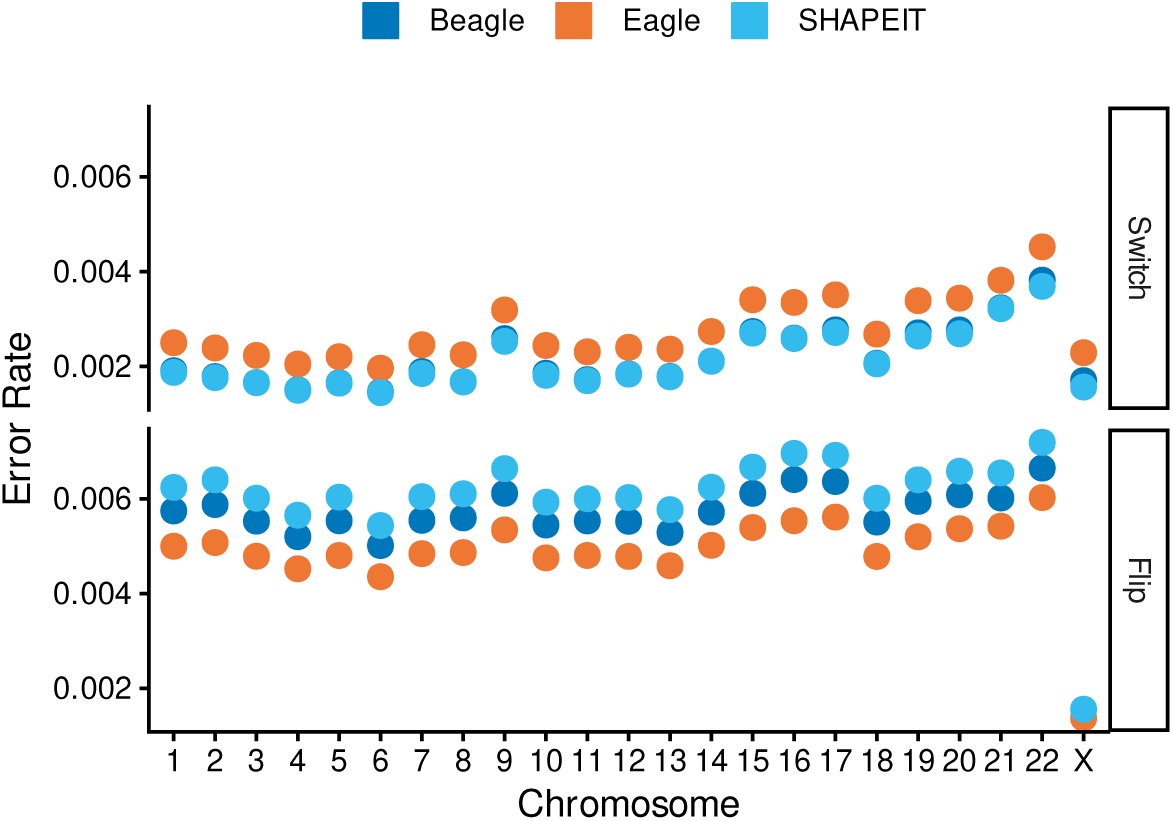
Average switch and flip rates across chromosomes. Each point represents the mean error rate (per heterozygous site) averaged across all samples for the autosome (trio rephasing) and chromosome X (synthetic diploids). **(A) Switch error rates:** Mean switch error rates. The average error rates observed in X for all three methods falls within the range of values observed in the autosomes. **(B) Flip error rates**: In X, we observe a notably smaller flip error rate than what is observed in the autosomes across all three methods.

The distribution of genomic predictors of phasing error also varied across chromosomes. Across the autosomes, the mean number of heterozygous sites ranged from 4.9 to 7.1 per 10 kb, higher than the 4.5 per 10 kb observed in the synthetic X-diploids (Supplementary Table B4). Similarly, the proportion of heterozygous sites at rare variants (maf *≤* 0.05) was slightly elevated in the autosomes (0.066–0.075) compared to the X chromosome (0.060). The representation of CpG sites at heterozygous positions was also generally higher on the autosomes, ranging from 0.142 to 0.212, relative to 0.142 on the X chromosome (Supplementary Figure A15). These genomic differences corresponded to divergent error enrichment profiles between X and the autosomes. While switch errors showed a consistent proportional enrichment at CpG sites across all chromosomes, the enrichment of flip errors at CpG sites was markedly lower on the autosomes (mean OR = 1.21) than on the X chromosome (mean OR = 2.92). In contrast, flip errors showed a much stronger association with rare variants on the autosomes, where odds ratios ranged from 33 to 149, substantially exceeding the range of 18 to 19 observed on the X chromosome (Supplementary Figures A21, A22). Enrichment of switch errors at rare variants, however, remained consistent across both the autosomes and the X chromosome.

## 3 Discussion

Statistical phasing enables downstream analyses such as imputation and the investigation of compound heterozygosity. However, it is challenging to evaluate the performance of statistical phasing for a specific dataset. While sequencing trios is typically used to generate high quality haplotypes, such trios are not always available and this practice can create biased phasing error rate estimates [9]. However, population studies are rich in haplotypes of known phase from male X chromosomes. Here in this study, we demonstrated the utility of synthetic diploid generated from male X chromosomes to directly benchmark the performance of statistical phasing.

We assessed the utility of this approach for three widely-used statistical phasing algorithms: Beagle 5.4, Eagle 2.4.1, and SHAPEIT 4 for reference-based phasing on sequenced genotypes from the 1000 Genomes Project. Overall, all three methods achieve high accuracy with statistically significant differences in performance. We identified a trade-off between error types: SHAPEIT 4 demonstrates the lowest switch error rate but yields the highest flip error rate while Eagle generates a higher switch error rate. This ranking of methods by both switch and flip rates is observed to be consistent across populations, affirming the robustness of our benchmark. High switch error rates may impact analyses depending on long-range linkage disequilibrium, such as identity-by-descent segment detection [20, 21]. In contrast, higher rates of flips may lead to less precise localized phase determination, which is critical for identifying compound heterozygous variants.

The synthetic diploids also allow for us to evaluate the genomic context where errors occur. We observed a significant enrichment of flip errors at rare variants (*maf ≤* 0.05) and CpG sites for all three methods. This is likely driven by the fact that rare variants are poorly tagged by common variation, and CpG sites are prone to parallel mutations on different haplotype backgrounds. The tendency of SHAPEIT4 to generate flip errors at low-frequency variants is well appreciated; while recent software updates include an imputation-like second stage for rare variants [3]. To approximate the impact of this extension, we reevaluated all phasing methods after removing variants with *maf <* 0.001 and observed that SHAPEIT4 still yielded a higher average flip rate than both Eagle and Beagle (Supplementary Figure A20). For switch errors, the strong positive correlation with recombination rate (*r ≈* 0.8) for SHAPEIT and Beagle highlights the intrinsic difficulty of maintaining phase across short LD blocks. Although Eagle’s switch error rate was higher overall, its correlation with recombination rate was lower (*r* = 0.45), suggesting that Eagle’s switch rate is driven by additional factors unrelated to the physical recombination rate.

Assessing the generalizability of these observations across the genome, our re-phasing of 602 trio probands confirmed that rankings of methods by switch and flip error rates remain consistent across the autosomes. Specifically, Eagle consistently exhibited the highest switch error rate, while SHAPEIT retained the highest flip error rate. While the rates of switch errors differ between the autosomes by up to 2-fold, the rates of all three methods on the X are comparable to autosomal rates, with X having very similar switch rates to chromosomes 1-8 and 9-13 (70% of the genome). However, we observed a marked difference in the number of flip errors, where we infer *>* 3*x* as many errors on autosomes compared to the X-chromosome. While autosomes carry more rare variants, this difference is insufficient to explain the increased number of errors. A likely contributor to this difference is the bias on phasing error rate estimates [9] created by genotyping errors as genotype errors both in parents and in offspring can lead to a perceived flip error. Even at the high quality of 30x genotyping, we expect 1 error per Mb (assuming 1 polymorphic site/1000 bp and an error rate of 0.001 [22] in each of the family members, a number that is higher than the rate of flip errors observed on the X chromosome (0.7 errors/Mb). Genotyping errors driving the estimated flip error rate would also explain the much weaker enrichment of CpGs among flip errors on the autosomes. As genotyping errors are not expected to accumulate at CpG positions, including genotyping errors in the flip errors waters down the enrichment of CpGs among the real flip errors. Overall, these results suggest that the synthetic diploid approach is an effective tool for benchmarking the relative performance of phasing algorithms across the genome, and may be more suitable to estimate the flip error rate than family-based benchmarks.

While our evaluation focused on reference-based phasing, similar considerations apply to population-based (*de novo)* phasing. In a *de novo* context, phasing on the X chromosome may also differ slightly from autosomal phasing due to the presence of perfectly phased males and a smaller number of haplotypes. Comparing reference-based phasing to *de nove* phasing, previous work [7] found similar switch error rates in samples phased both with and without a reference panel, suggesting the comparative performance and genomic context of errors identified here likely persist across different phasing modes. By characterizing both the frequency and genomic context of errors, this work provides evidence-based guidance for researchers selecting appropriate software and interpreting analyses sensitive to haplotype length or local phase.

Overall, the proposed method of using synthetic X-diploids enables detailed characterization of phasing errors for reference-based phasing with the three most commonly used methods. Our findings remain consistent across diverse populations and are broadly generalizable to autosomal estimates, suggesting that the underlying drivers of phasing difficulty are largely universal. This approach can be easily implemented in any population-scale study that includes male probands: synthetic X-diploids can be phased alongside the study samples to provide immediate, dataset-specific insights into algorithmic performance.

## 4 Methods

We used male X chromosomes to construct synthetic diploids of known phase in order to evaluate three model statistical phasing algorithms [1–3]. The chromosomes were sampled from the 2,504 phase 3 samples of the 1000 Genomes Project (1kGP), allowing for comparisons to be made across distinct ancestral groups. In addition to calculating error rates, we also characterized the genomic context in which errors occur by comparing error densities across the chromosome to other genomic features. Additionally we rephased the autosomes of the probands from 602 trio pedigrees and evaluated the discordance between the population-based phasing inferred haplotypes and the pedigree-adjusted phase.

### 4.1 1000 Genomes 30x High-Coverage Sequencing Data

The high-coverage WGS expanded 1kGP cohort consists of the 2,504 original phase 3 1kGP samples (1233 males and 1271 females), along with 602 trios with probands having had their phase refined using Mendelian inheritance logic. All of these samples are members of one of five super-populations (AFR, AMR, EAS, EUR, SAS), with each super-population consisting of non-overlapping subsets of 26 distinct populations [12]. For the construction of the synthetic chromosome X diploids we used only the 2,504 phase 3 unrelated individuals. From the phased VCF for chromosome X we removed sites that are in either pseudo-autosomal region (PAR1 and PAR2), and we also remove sites that are difficult to sequence according to the 1kGP pilot accessibility mask [23]. Non-SNP variants were also removed, as were SNPs who were either non-biallelic or singletons.

For our analysis of the 602 1kGP trios, we applied the same criteria, filtering the phased and unphased VCF files for all autosomes to retain only biallelic, non-singleon SNPs that are not in chromosomal regions covered by the 1kGP pilot accessibility mask.

### 4.2 Synthetic Diploid Construction and Phasing

To create synthetic diploid individuals for benchmarking, we first defined the sampling pool by selecting a population from a given 1kGP super-population. From this pool, we sampled pairs of male X chromosomes and combined them to form a synthetic diploid, ensuring that no pair of samples was chosen more than once. Based on the sampled haplotypes, we created 2 files: a phased VCF file with the known phase and an unphased VCF where at heterozygous sites the order of the two alleles is selected at random. For the phasing step, we created a unique reference panel for each synthetic diploid, consisting of the remaining 2,502 samples not used in its construction. In total, we constructed 200 synthetic diploids for each of the five 1kGP super-populations in this manner. Each synthetic diploid was then phased using Eagle v2.4.1, Beagle 5.4, and SHAPEIT4 using default options and genetic maps provided by the respective authors [1–3].

### 4.3 Trio Proband Phasing

For each of the 602 1kGP trios, we rephased the autosomes of each proband without pedigree-based correction. Each proband was phased individually using a reference panel constructed from the 2,504 unrelated 1kGP phase III samples, excluding all family members. To avoid potential errors, we excluded triple heterozygous positions at which patterns of Mendelian inheritance cannot be used to resolve haplotypes using only a trio. We also exclude these positions from the original, trio-resolved haplotypes phased with pedigree-based correction [12] against which we benchmarked the phasing algorithms.

### 4.4 Evaluation of Population-based Inferred Phase

For our synthetic X diploids and the rephased trios, we identified the location of switch and flip errors by comparing the inferred diploids from each of the three phasing methods to the original haploids used in construction of the synthetic diploid. Using Vcftools (version 0.1.17), we identified the location of any discordance between the inferred phase and the known true phase. Each discordance was then classified as one of two error types: flips, defined as a double-switch error where two consecutive phase errors occur at consecutive heterozygous positions; and switches, any non-flip discordance.

### 4.5 Genomic Predictors of Error

To assess the influence of specific genomic features on phasing accuracy, we evaluated the enrichment of errors at CpG sites, rare variants, and regions of high recombination. We identified CpG sites based on annotations in the genome assembly GRCh38. Minor Allele Frequencies (maf) were determined from the 1kGP deep sequence dataset used to construct the synthetic diploids. To quantify the relationship between phasing error and recombination rate, we used a high-resolution recombination map of the X chromosome derived from the 1000 Genomes Project data [23]. We divided the X chromosome into non-overlapping 1 MB bins. We then computed the average error rate (switches and flips separately) and the average recombination rate (weighted by the size of the original map intervals within the bin) for each 1 MB bin. Pearson correlation coefficients were then computed using these binned values to assess the association between error rates and the local recombination environment. Furthermore, to examine enrichment, we assigned each heterozygous site to a recombination rate decile based on the original map data. Finally, we used the Cochran-Mantel-Haenszel (CMH) test to evaluate the statistical significance of the enrichment of both switch and flip errors at CpG sites and within maf categories, treating each synthetic diploid as a stratum.

## Acknowledgments

This work was supported by NIH grant R01HG011031.

## Declaration of interests

The authors declare no competing interests.

## Data and code availability

The genomic data analyzed in this study are publicly available from the 1000 Genomes Project (1kGP). All code used for simulations, haplotype phasing benchmarks, and statistical analyses is available in the following GitHub repository: https://github.com/theandyb/phasing clean.

## Appendix A Supplementary Figures

**Fig. A1:**
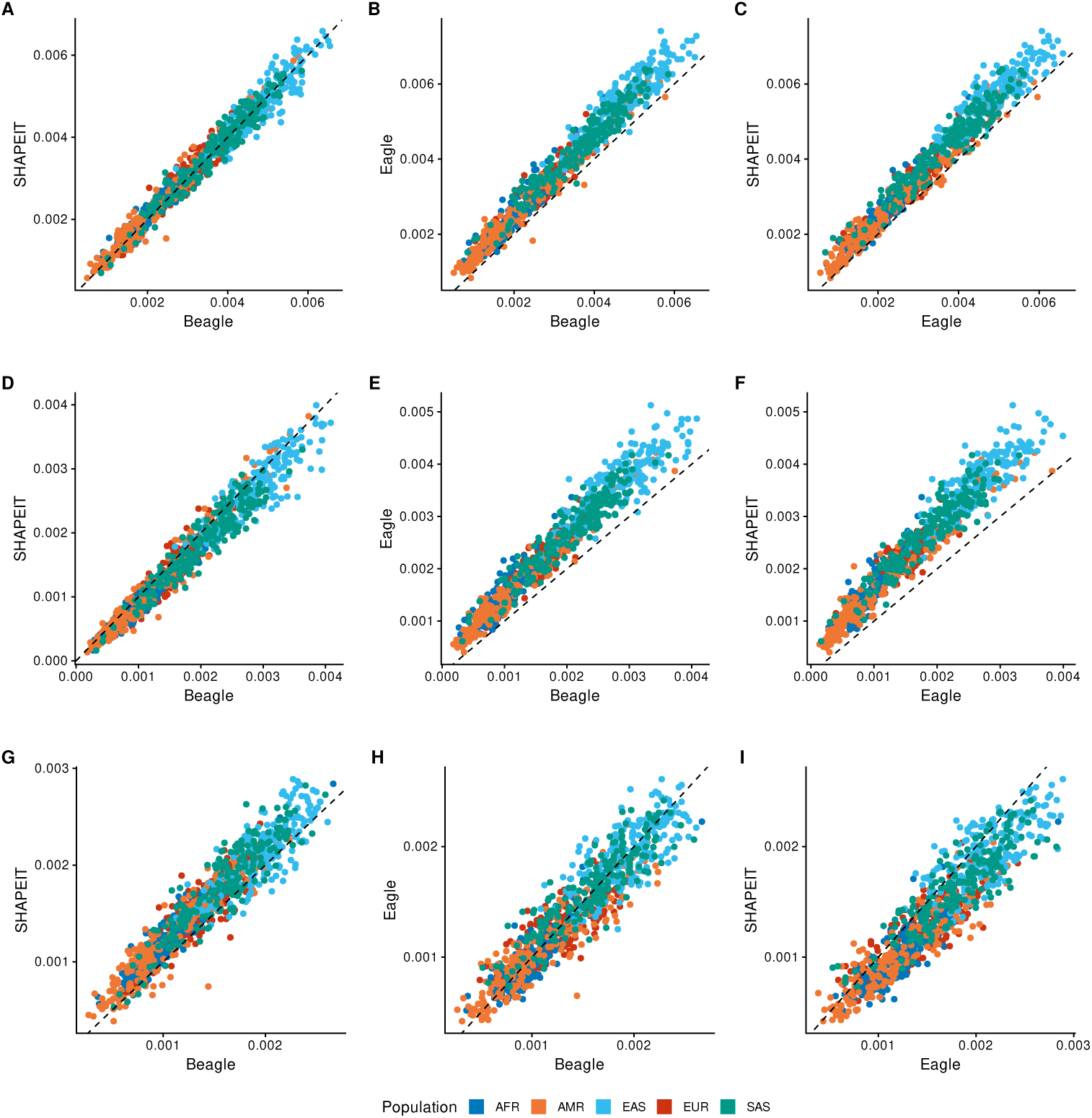
Error rates observed in each synthetic X chromosome diploid across each pair of methods. Rates are computed as the total number of errors divided by the number of heterozygous positions in each synthetic diploid. Each point represents one synthetic diploid with the x– and y-position determined by the total number of errors observed in each method when comparing the inferred haplotypes to the original male X chromosome haplotypes sampled to generate the synthetic diploid.

**Fig. A2:**
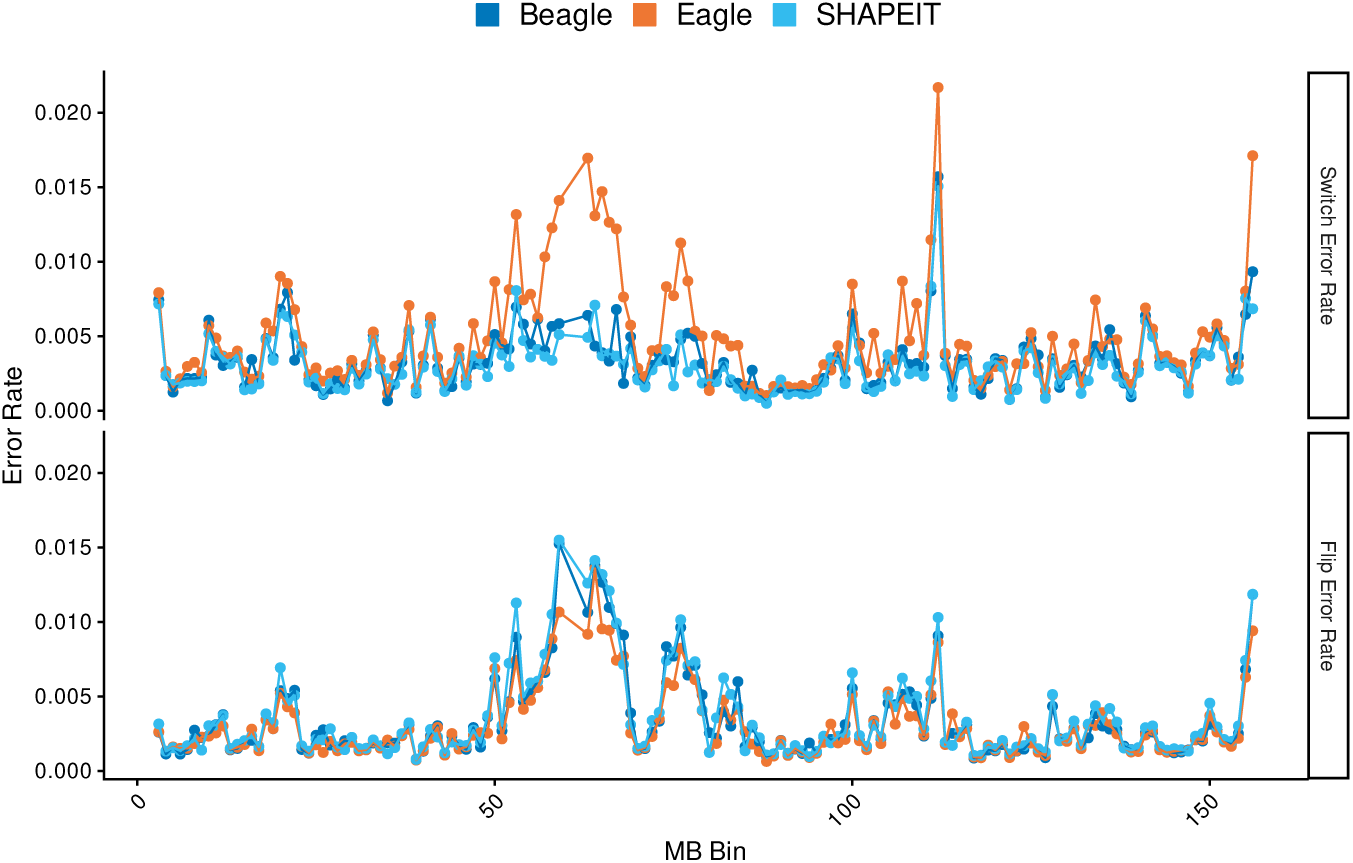
Average switch and flip error rates are shown in 1-Mb non-overlapping windows across the X chromosome. Each point on the line represents the mean error rate for all 1,000 synthetic diploids within that specific genomic bin. For both switch and flip errors, the rates for each method are highly correlated across the chromosome. However, for switch errors, Eagle (yellow line) exhibits several localized spikes where its error rate is markedly elevated compared to both Beagle and SHAPEIT.

**Fig. A3:**
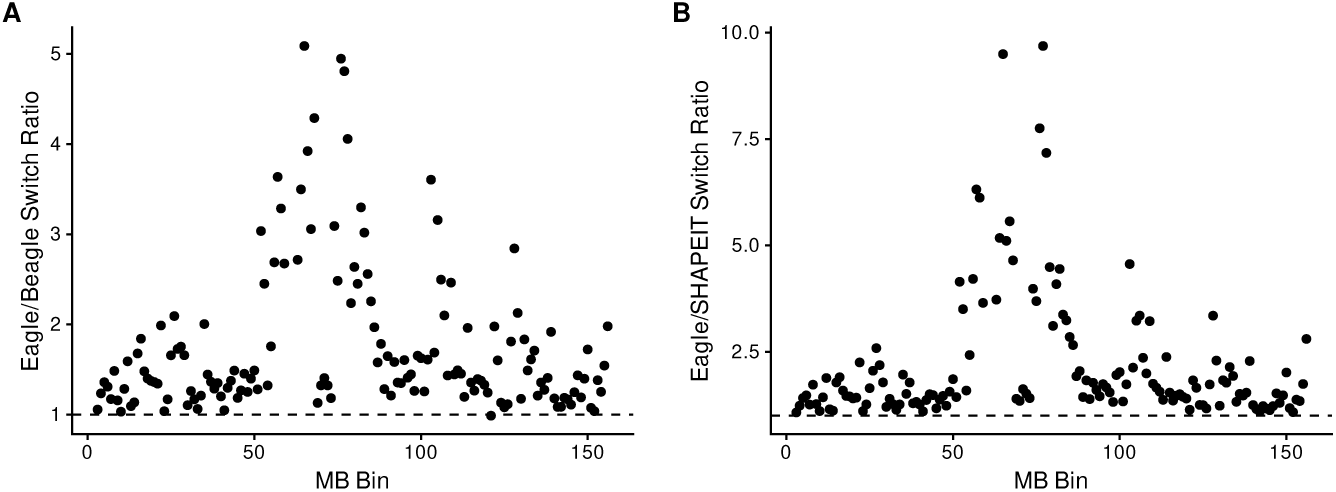
Enrichment of Eagle switch error rates in MB bins relative to Eagle and SHAPEIT. **(A)** Ratio of Eagle switch error rates to Beagle. On average each bin’s average switch rate for Eagle was 1.75 times that of Beagle. **(B)** Ratio of Eagle switch error rates to SHAPEIT. On average each bin’s average switch rate for Eagle was 2.18 times that of Beagle.

**Fig. A4:**
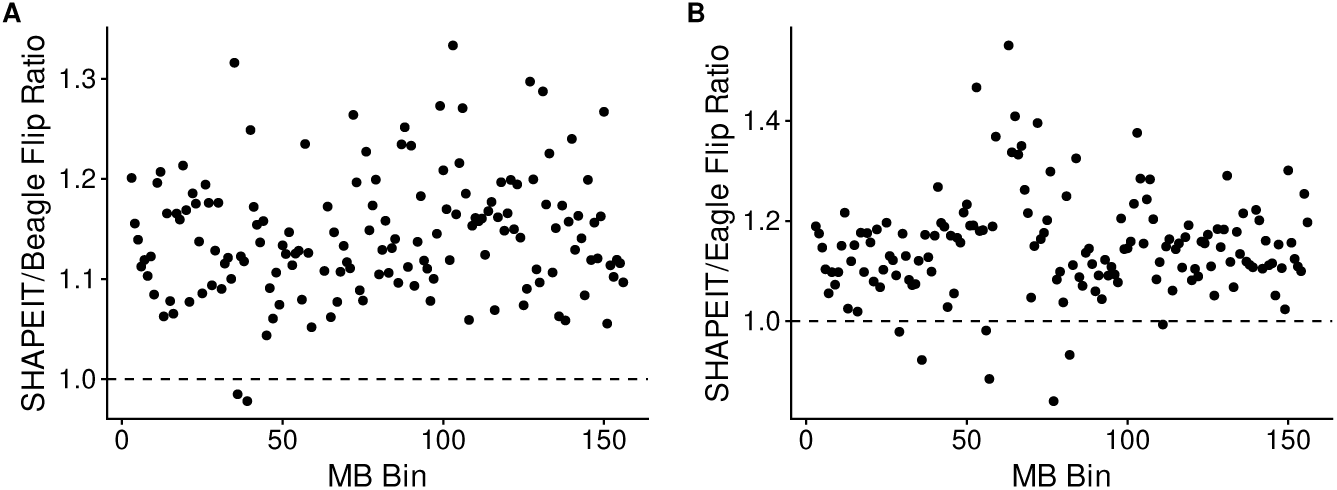
Enrichment of SHAPEIT flip error rates in MB bins relative to Beagle and Eagle. **(A)** Ratio of SHAPEIT flip error rates to Beagle. On average each bin’s average flip rate for SHAPEIT was 1.14 times that of Beagle. **(B)** Ratio of SHAPEIT switch error rates to Eagle. On average each bin’s average switch rate for SHAPEIT was 1.15 times that of Eagle.

**Fig. A5:**
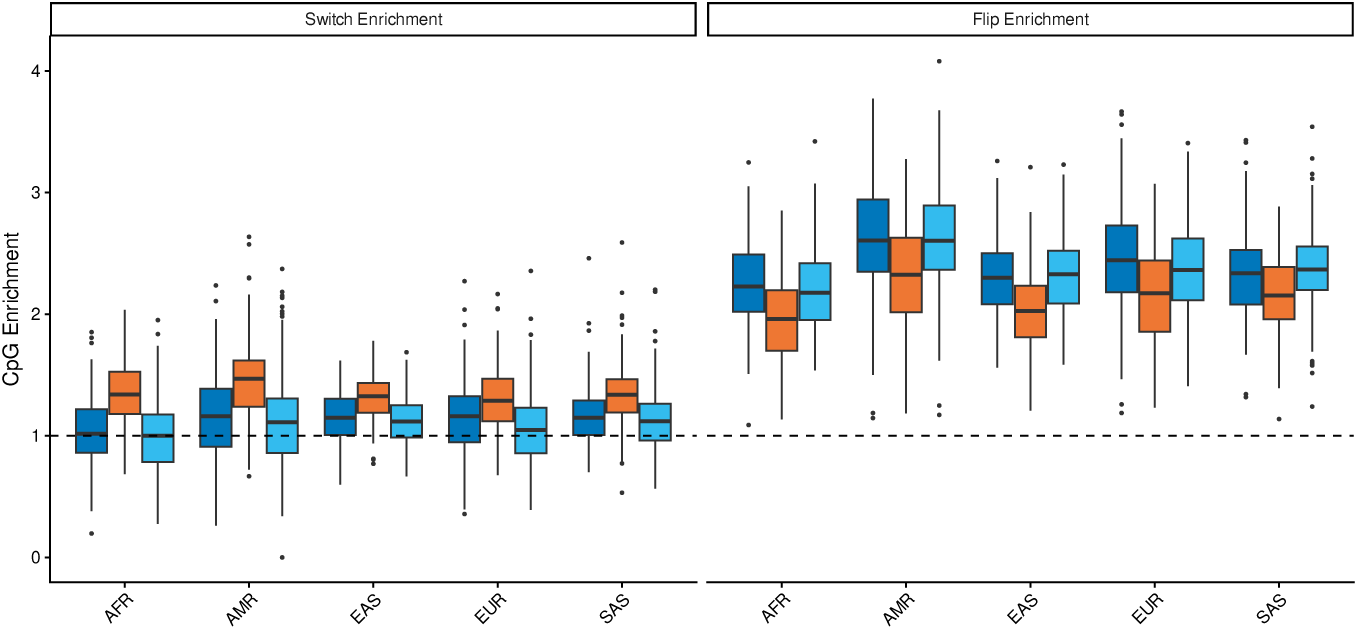
Distributions of switch error and flip error enrichment observed in chromosome X synthetic diploids for each method by population. For each sample, enrichment is computed by dividing the fraction of errors occurring at CpG sites by the fraction of heterozygous sites at CpG sites. Across methods, both switch (*F* (8, 1990) = 7.49*, p <* 6.88 *×* 10*^−^*^10^) and flip (*F* (8, 1990) = 4.23*, p <* 4.76 *×* 10*^−^*^5^) error rates were found to vary across populations.

**Fig. A6:**
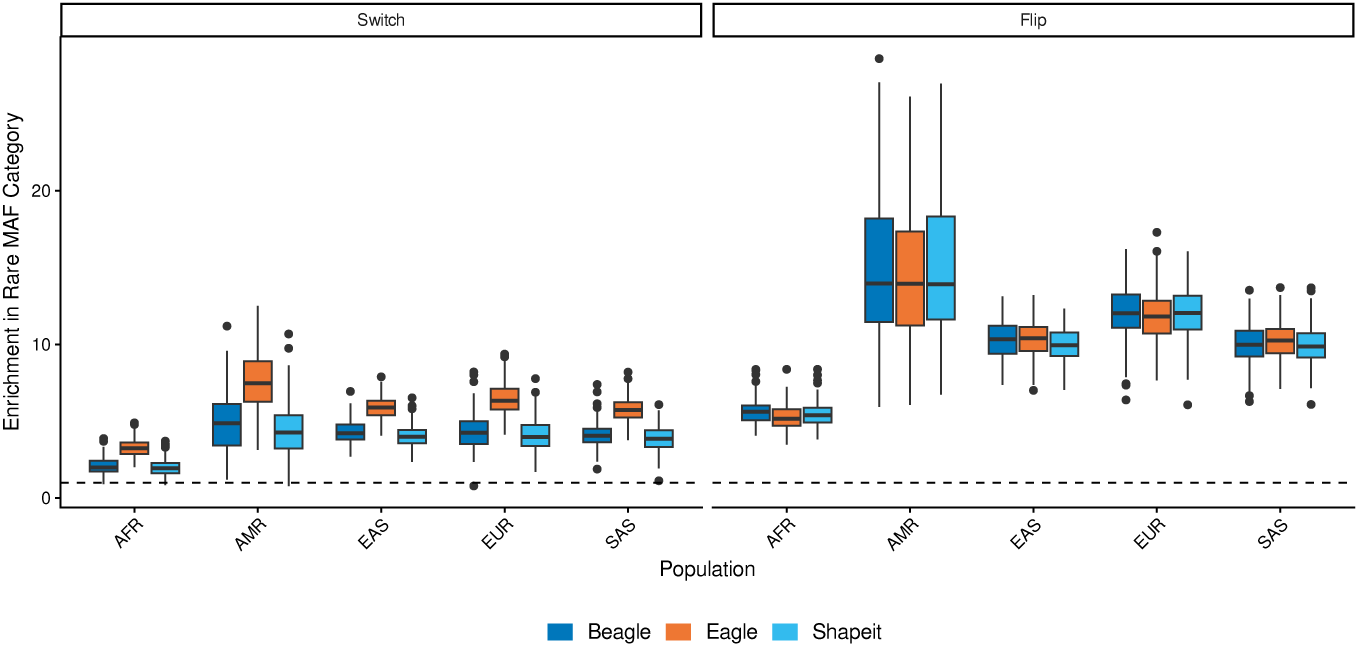
Enrichment of phasing errors at rare variants by population. For each sample, enrichment is computed by dividing the fraction of errors occurring at rare (*maf <* 0.05) variants by the fraction of heterozygous sites at rare variants. Across methods, both switch (*F* (8, 1990) = 51.42*, p <* 2.2 *×* 10*^−^*^16^) and flip (*F* (8, 1990) = 13.92*, p <* 2.2 *×* 10*^−^*^16^) error rates were found to vary across populations.

**Fig. A7:**
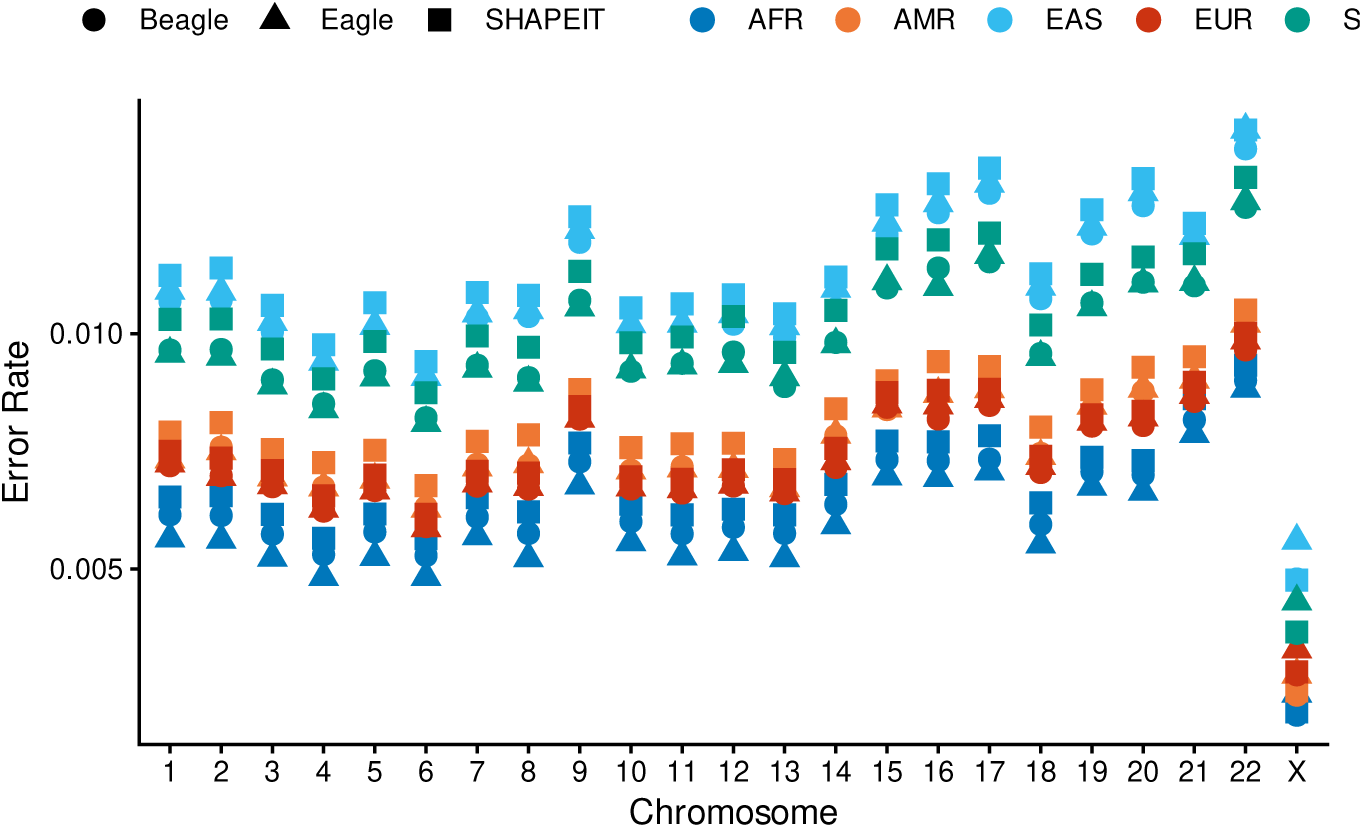
Phasing error rates observed across the genome, comparing trio-derived autosomal error rates to synthetic X-diploid rates. Phasing method is indicated by point shape, while 1kGP populations are indicated by color.

**Fig. A8:**
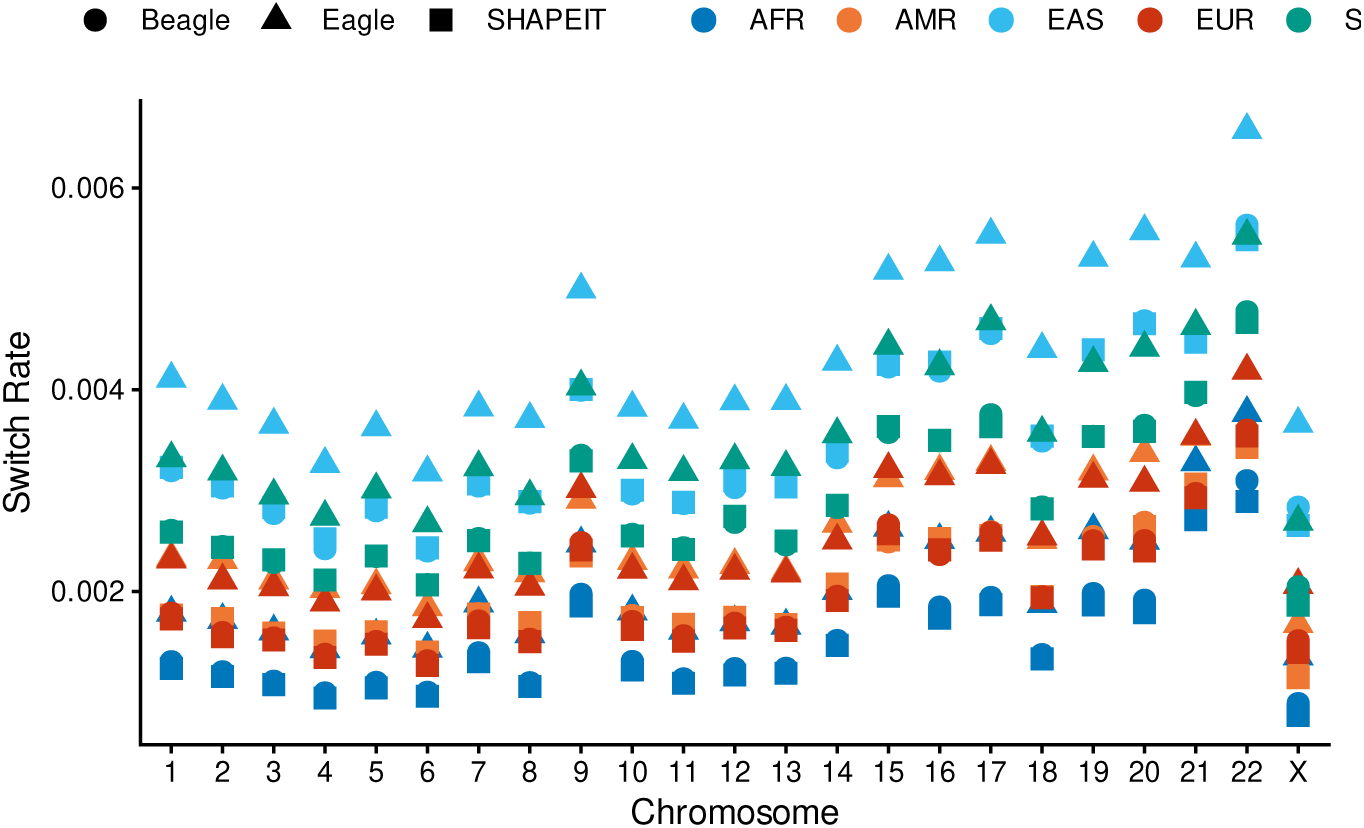
Mean switch error rates (single switch events) per chromosome. Phasing method is indicated by point shape, while 1kGP populations are indicated by color. Autosomal rates reflect trio proband re-phasing without parental context, provided for comparison against synthetic X-diploid results.

**Fig. A9:**
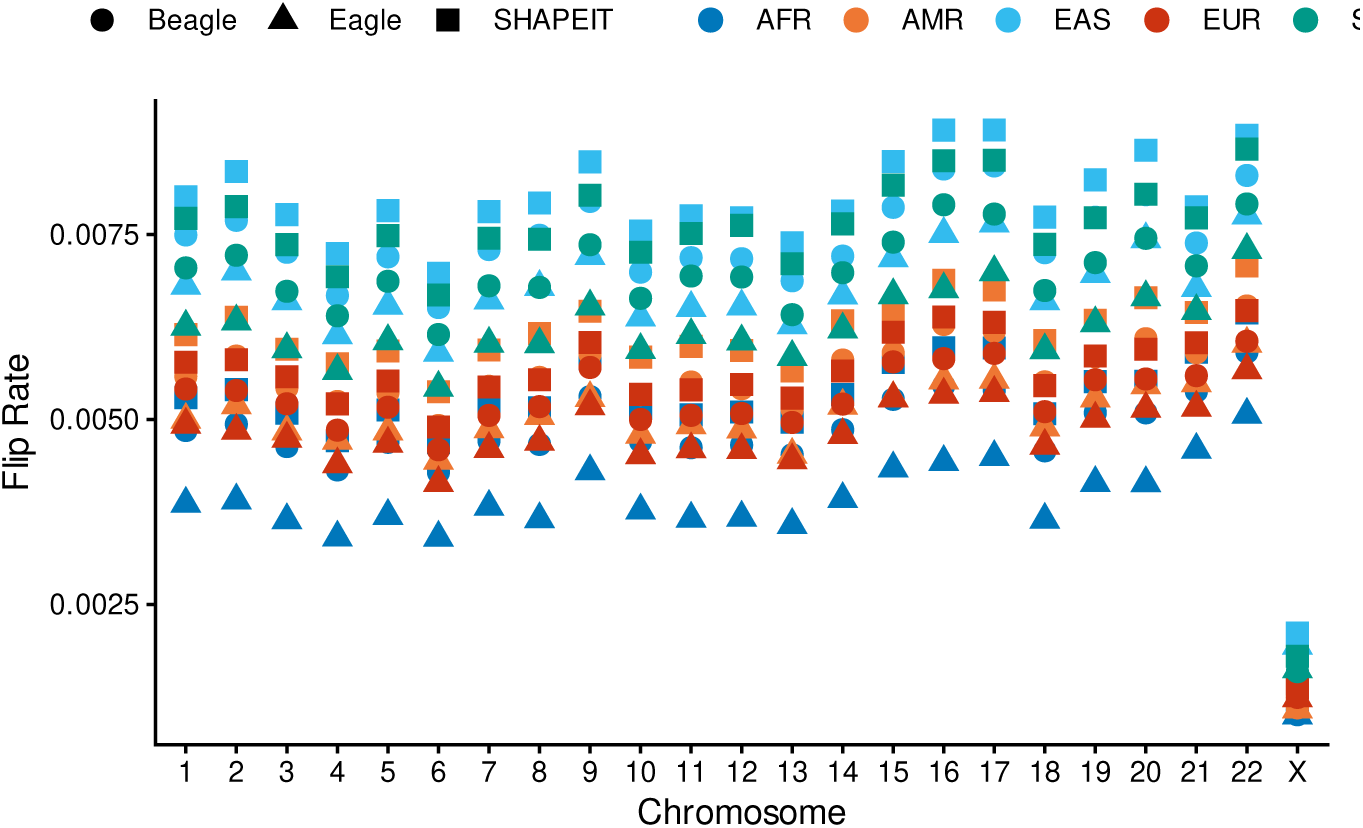
Mean flip error rates (double switch events) per chromosome. Phasing method is indicated by point shape, while 1kGP populations are indicated by color. Autosomal rates reflect trio proband re-phasing without parental context, provided for comparison against synthetic X-diploid results.

**Fig. A10:**
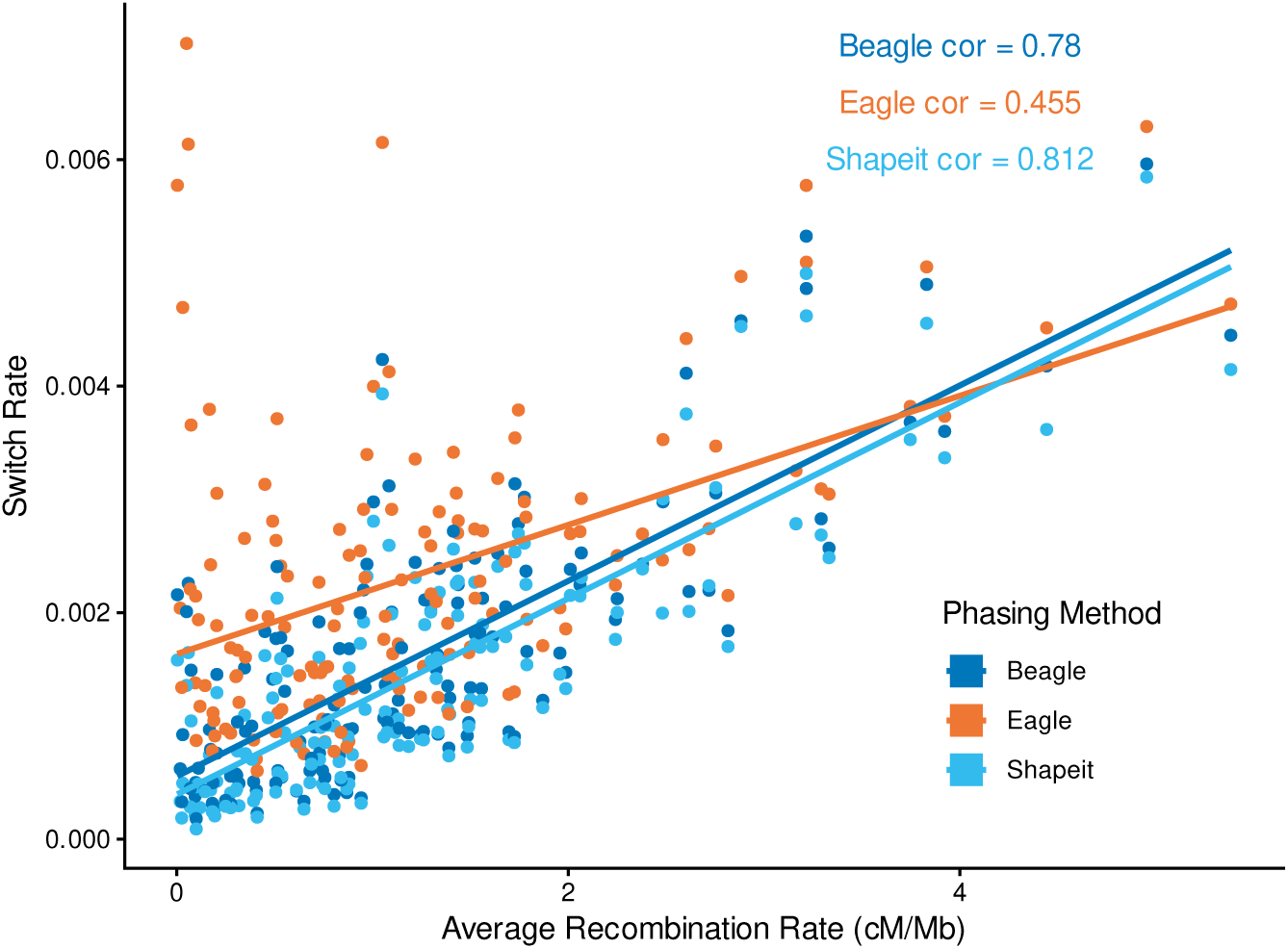
Average switch error rates and recombination rate values in non-overlapping MB bins in chromosome X. Switch error rates are observed to be correlated with average recombination rates in non-overlapping MB bins in chromosome X (Beagle: *r* = 0.78*, p <* 0.001; Eagle: *r* = 0.45(*p <* 0.001); SHAPEIT: *r* = 0.81(*p <* 0.001)).

**Fig. A11:**
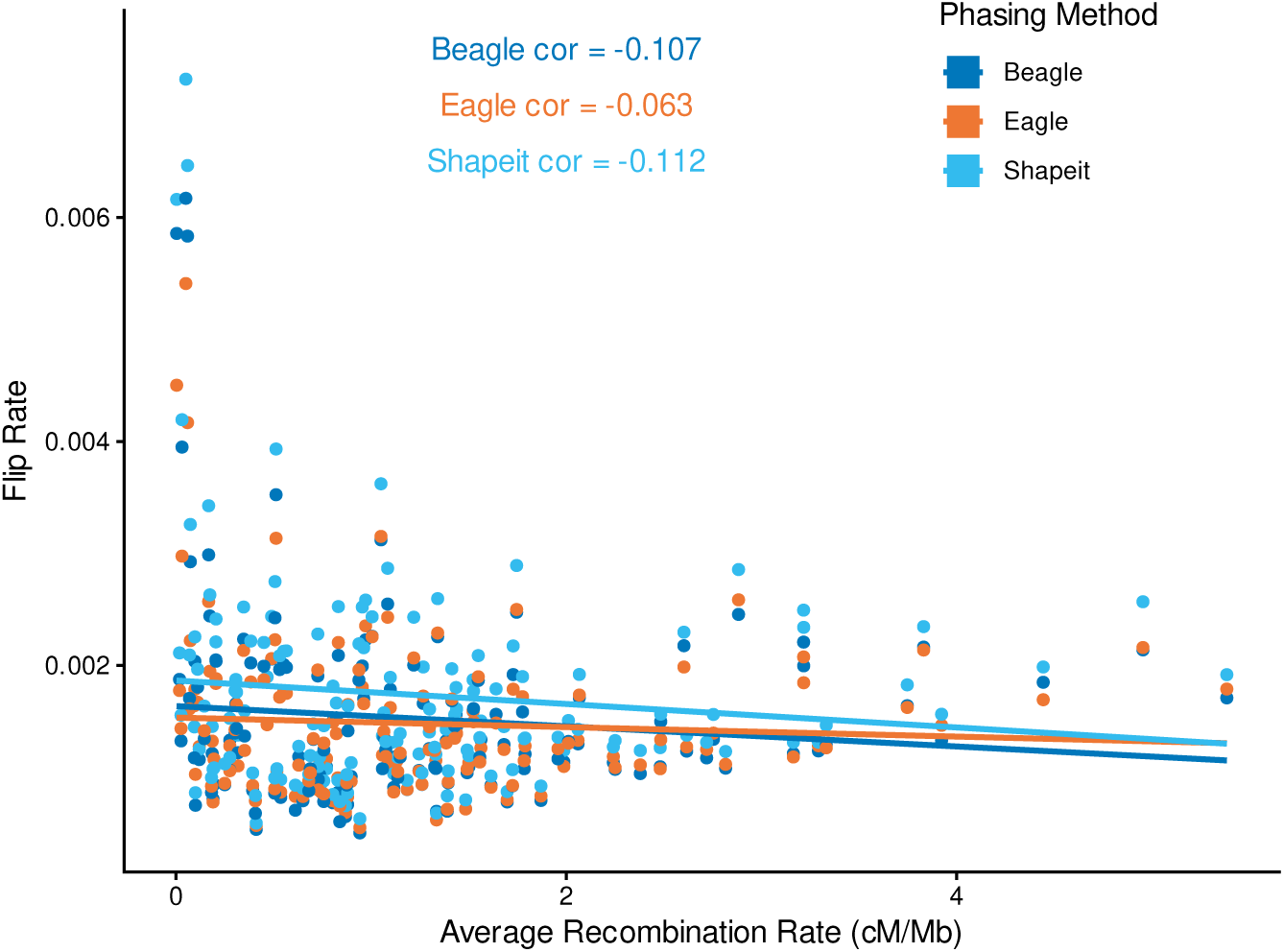
Average flip error rates and recombination rate values in non-overlapping MB bins in chromosome X. Flip errors are not found to be significantly correlated with recombination rates.

**Fig. A12:**
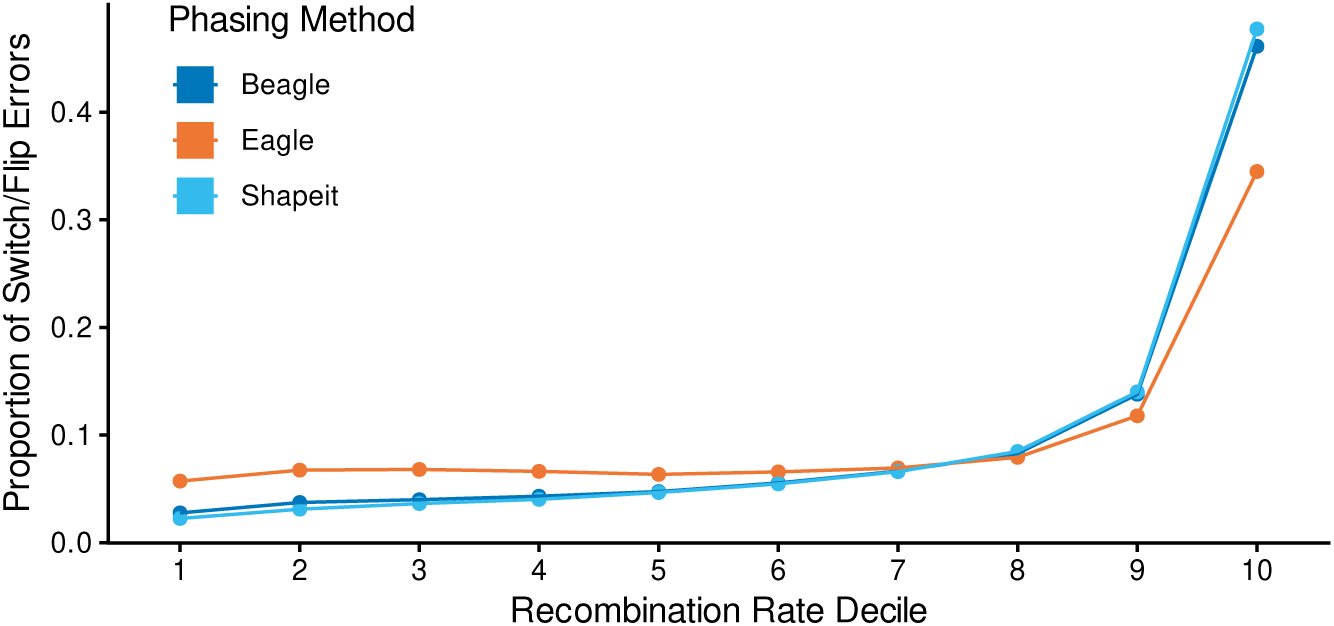
Proportion of switch errors stratified by recombination rate deciles. Mean proportion of total switch errors occurring at heterozygous sites within each recombination decile. Results are averaged over 1000 synthetic diploids. The top decile (highest recombination rate) contains an enriched proportion of total switch errors (*>* 10%).

**Fig. A13:**
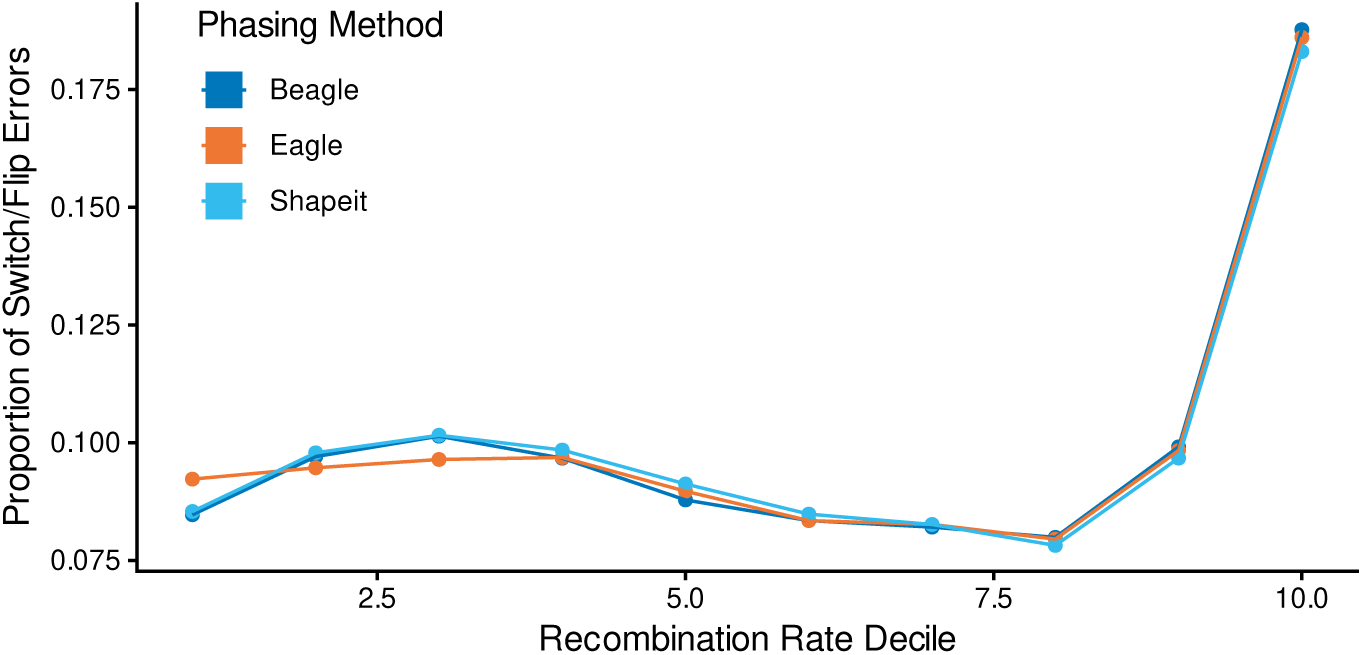
Proportion of flip errors stratified by recombination rate deciles. Mean proportion of total flip errors occurring at heterozygous sites within each recombination decile. Results are averaged over 1000 synthetic diploids. The top decile (highest recombination rate) contains an enriched proportion of flip errors (*>* 10%).

**Fig. A14:**
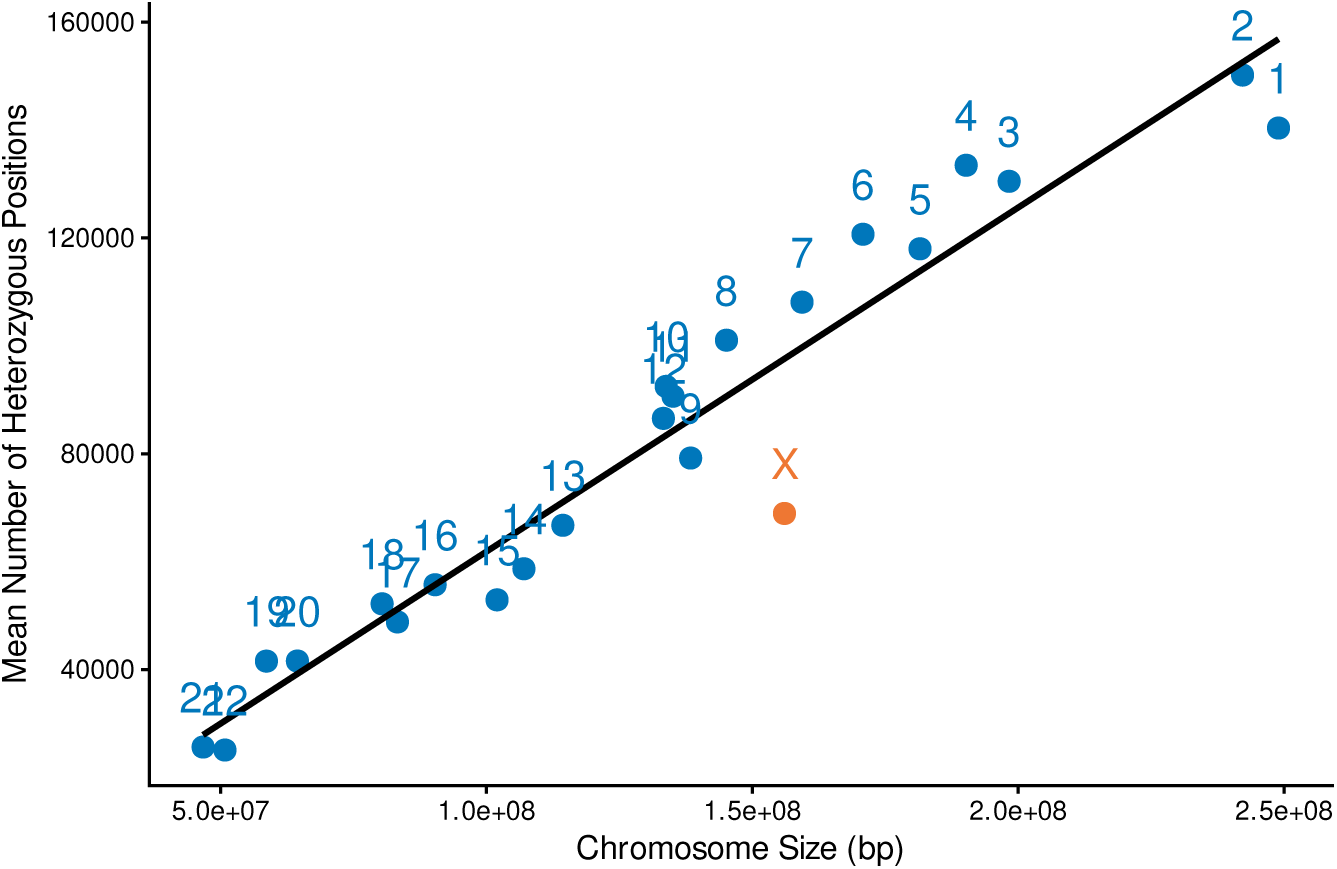
Relationship between chromosome size and heterozygosity. Mean number of heterozygous sites plotted against physical chromosome size (Mb) for autosomal trio probands and synthetic diploid X-chromosomes. The average number of heterozygous positions in chromosome X synthetic diploids is found to be less than similarly sized autosomes.

**Fig. A15:**
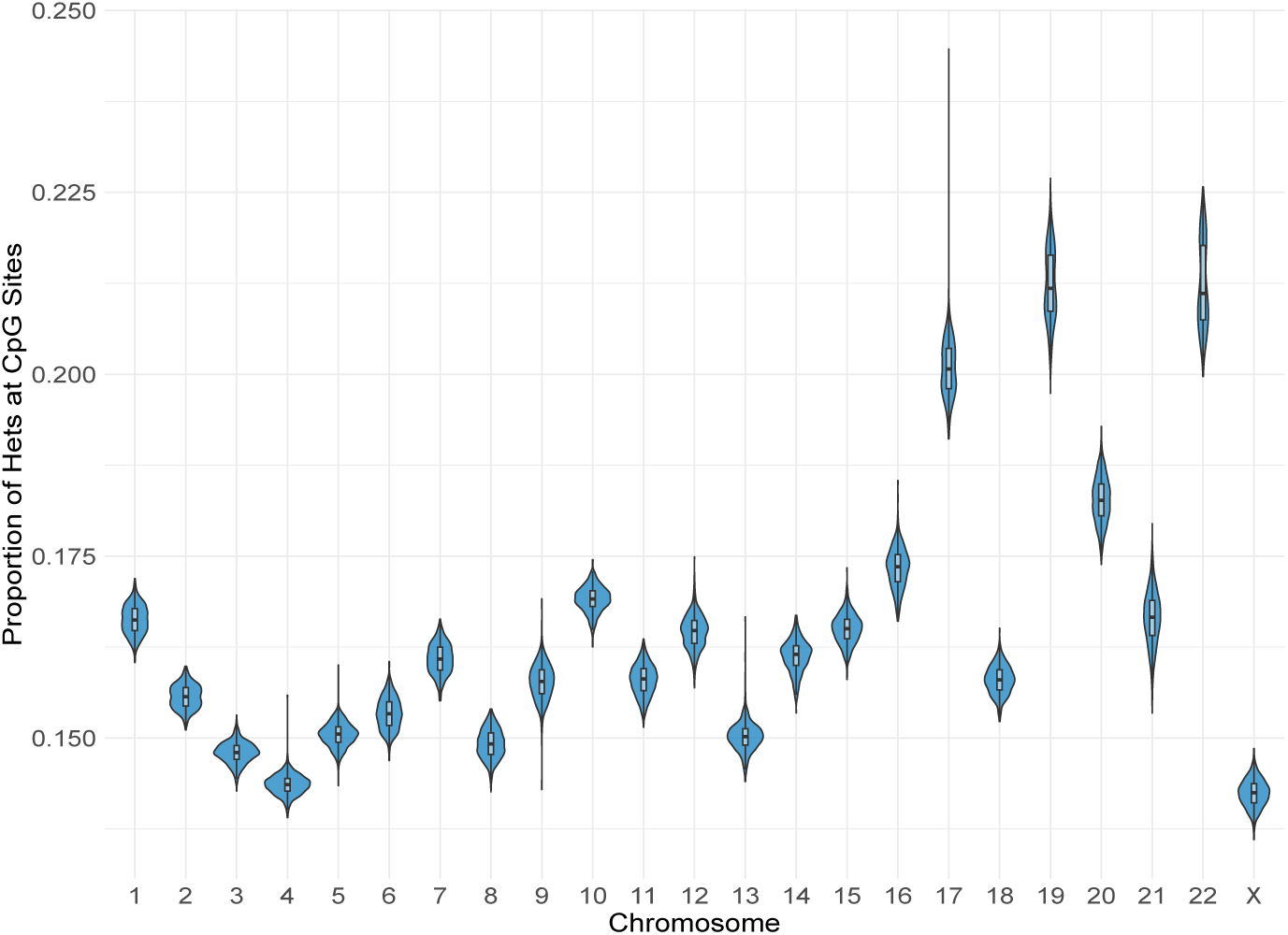
Distribution of the proportion of heterozygous sites occurring at CpG dinucleotides across all autosomes and chromosome X. Synthetic X-chromosomes show a reduced proportion of CpG-associated heterozygosity relative to the autosomes.

**Fig. A16:**
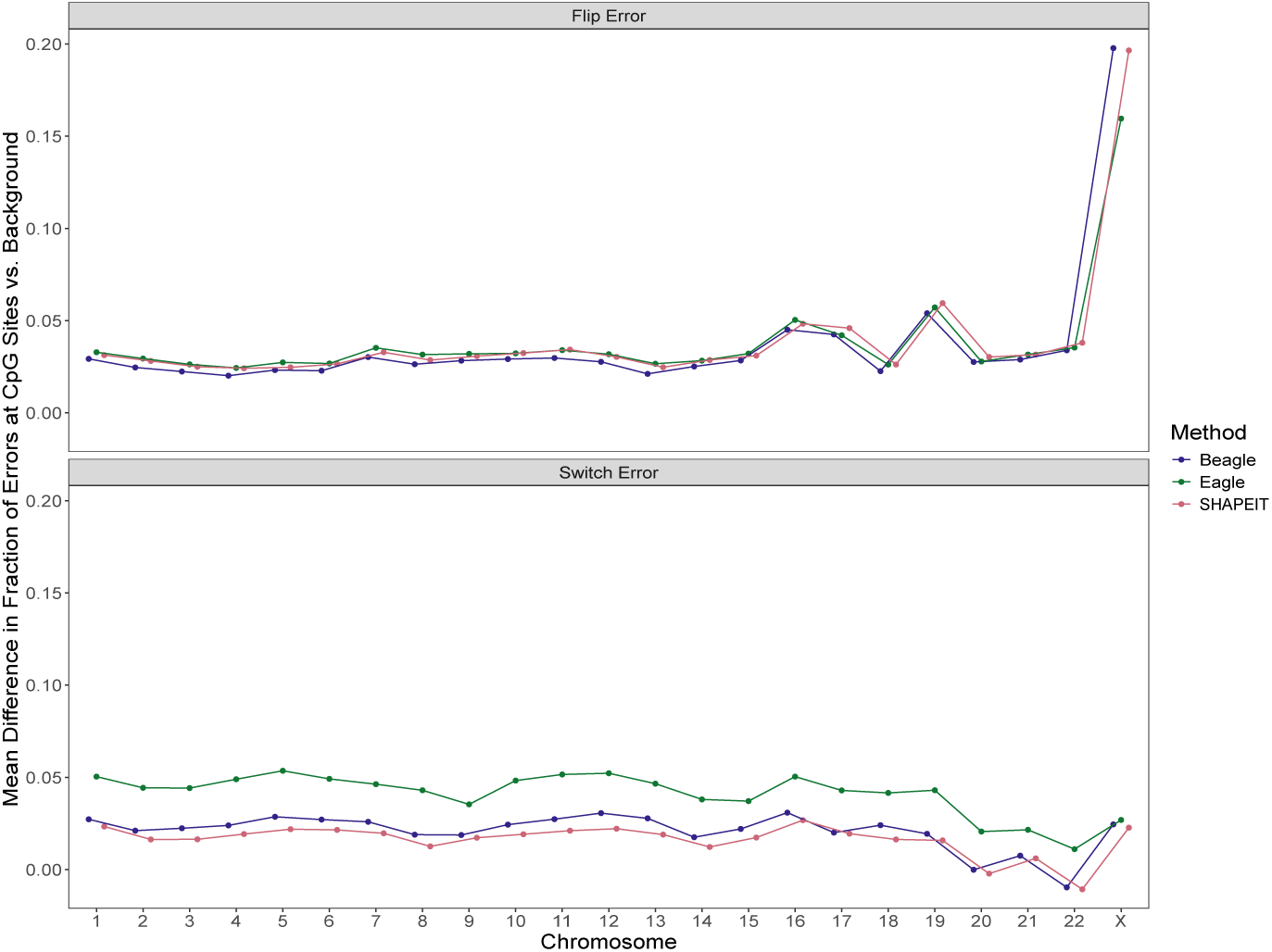
Enrichment of flips and switch errors at CpG sites across the autosomes (trio re-phase) and X (synthetic diploids). Enrichment is computed as the mean difference between the proportion of errors at CpG sites and the proportion of heterozygous positions at CpG sites.

**Fig. A17:**
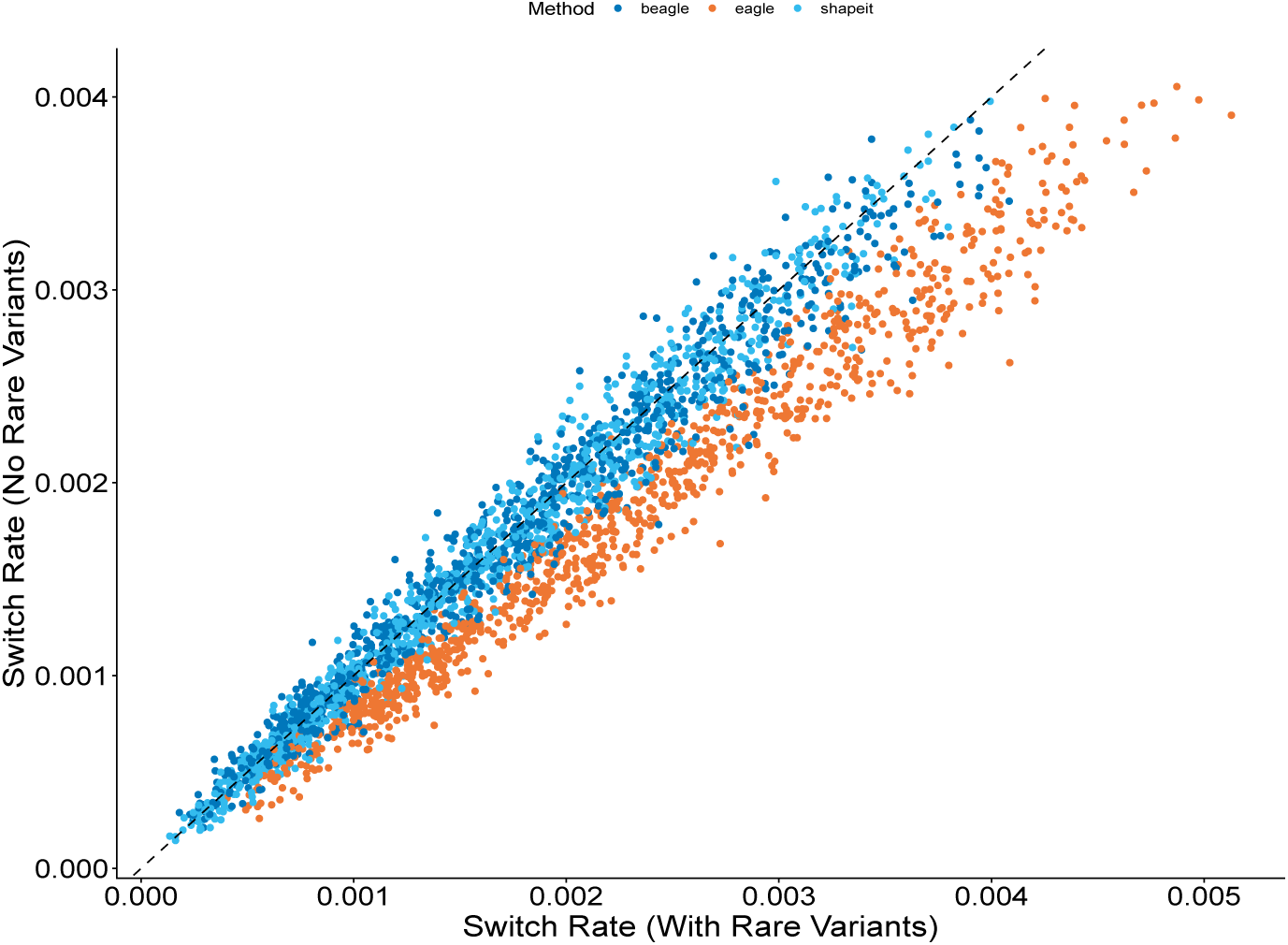
Switch rates observed in the 1000 synthetic diploids with and without very rare (*maf <* 0.001) variants.

**Fig. A18:**
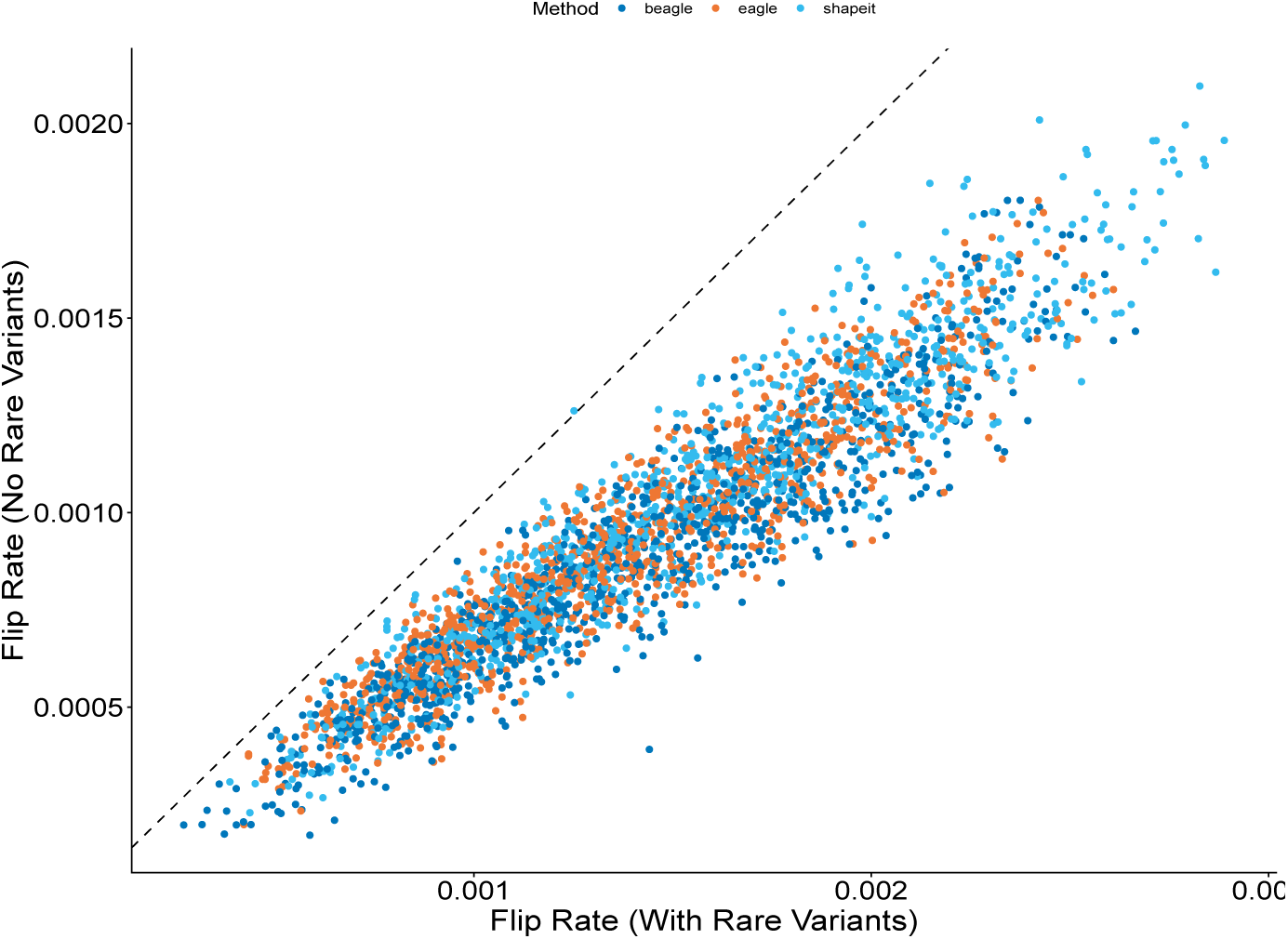
Switch rates observed in the 1000 synthetic diploids with and without very rare (*maf <* 0.001) variants.

**Fig. A19:**
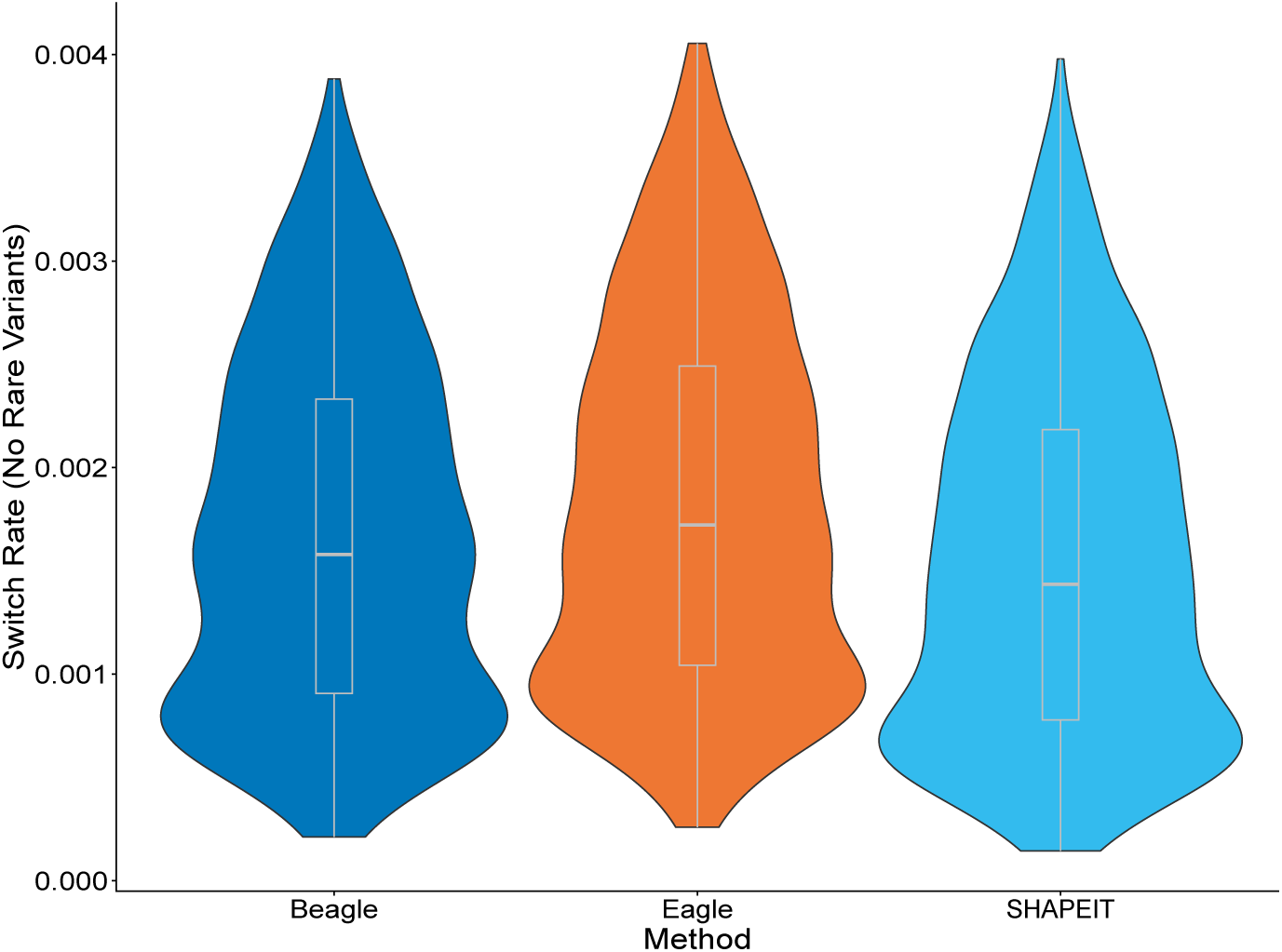
Distributions of switch error rates by method in the 1000 synthetic X diploids phased with very rare (*maf <* 0.001) variants removed.

**Fig. A20:**
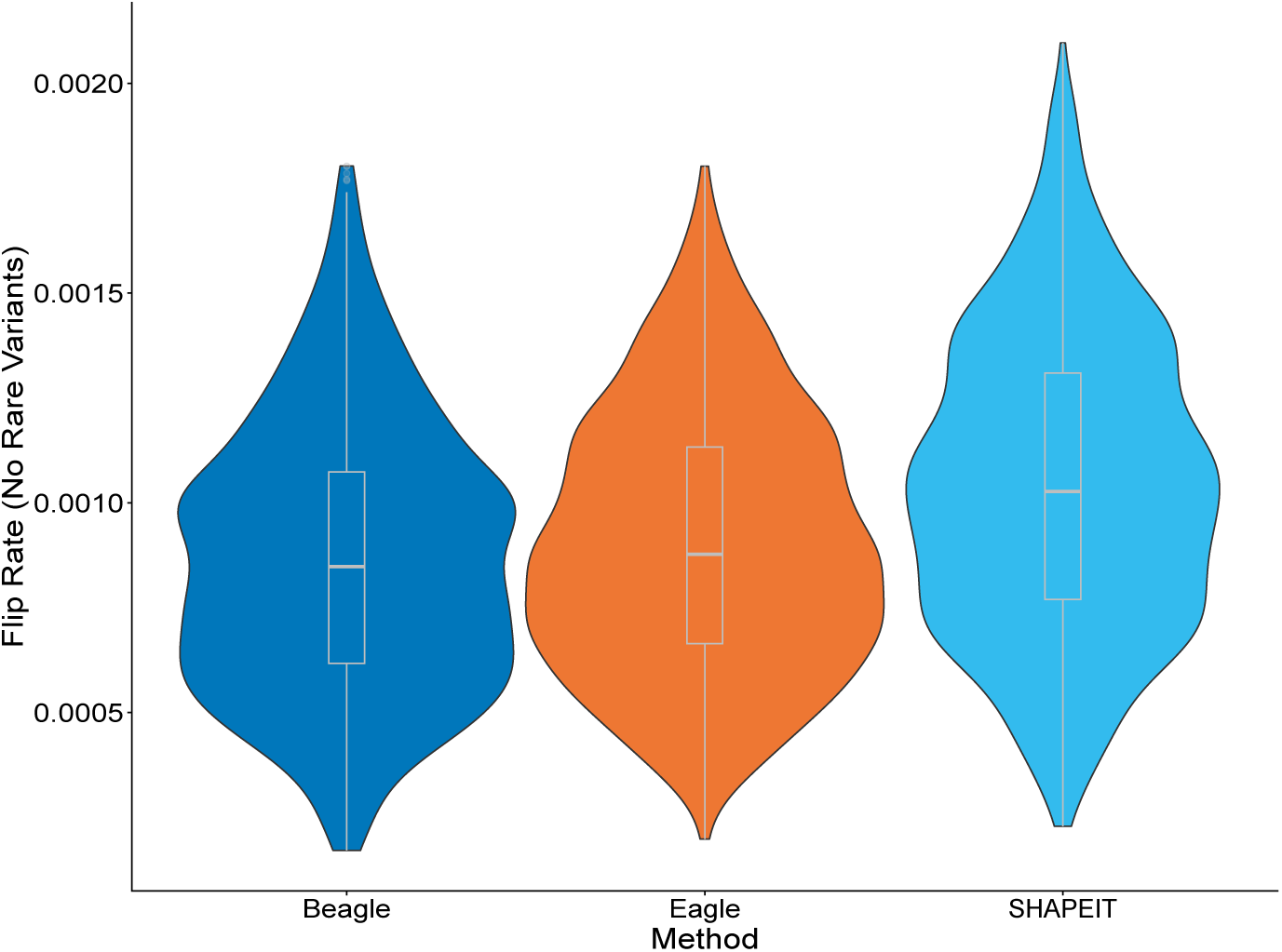
Distributions of flip error rates by method in the 1000 synthetic X diploids phased with very rare (*maf <* 0.001) variants removed.

**Fig. A21:**
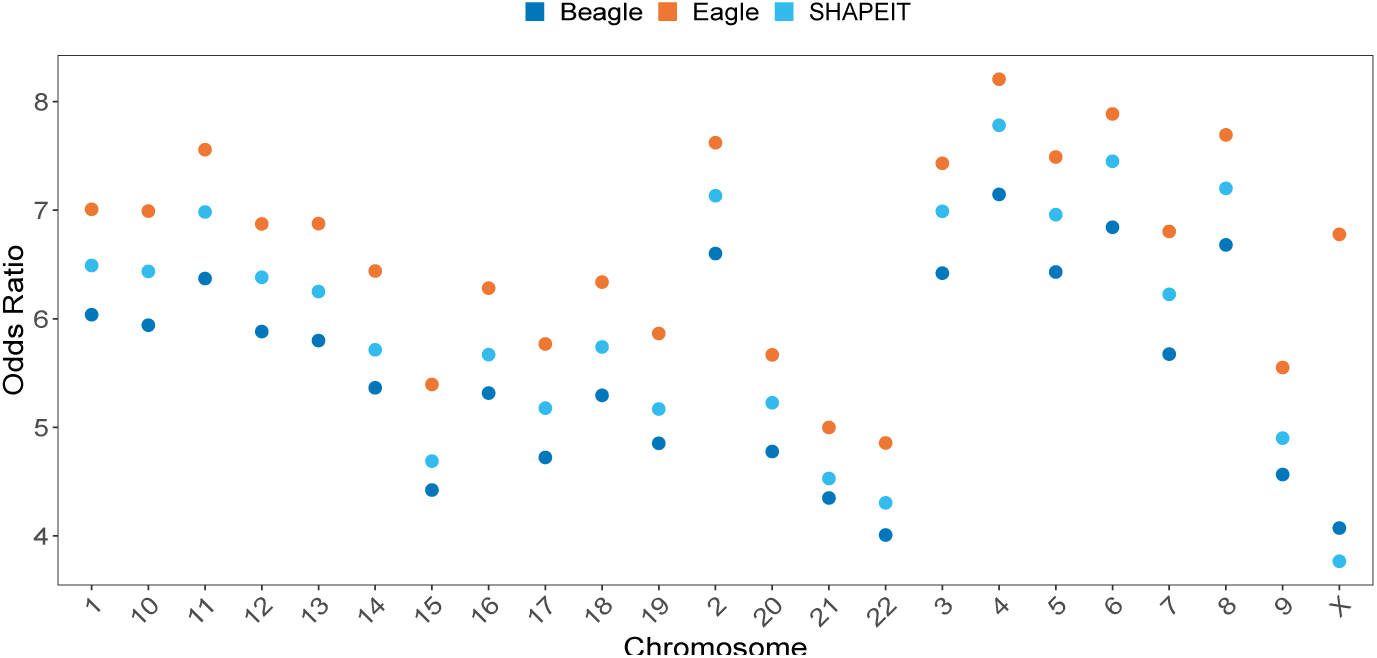
Odds ratios for switches at rare variants by chromosome and method. Data points represent the ratio calculated from the sum of all observations across all samples (*n* = 602 trio autosomes, *n* = 1000 X synthetic diploids).

**Fig. A22:**
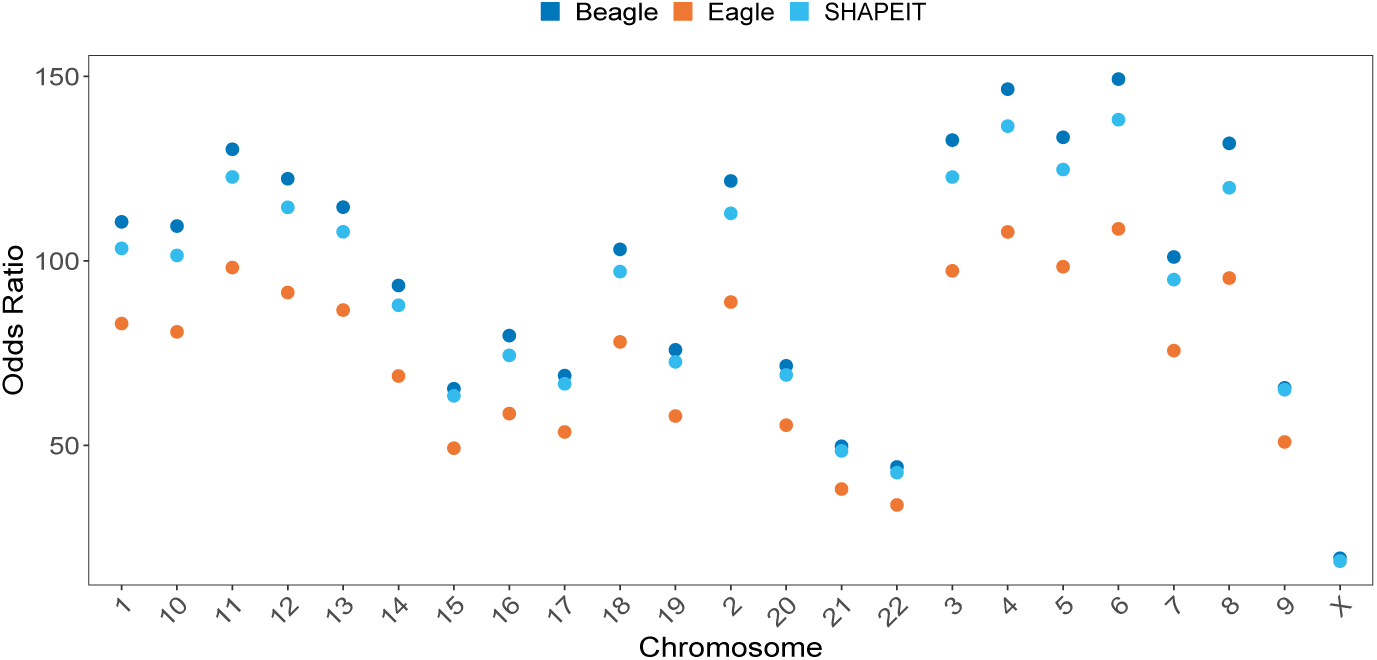
Odds ratios for flips at rare variants by chromosome and method. Data points represent the ratio calculated from the sum of all observations across all samples (*n* = 602 trio autosomes, *n* = 1000 X synthetic diploids).

## Appendix B Supplementary Tables

**Table B1:** T test results comparing the proportion of errors at CpG sites to the background proportion of heterozygous sites at CpG sites.

| Error Type | Method | Mean Difference | T | P value | CI Lower | CI Upper |
| --- | --- | --- | --- | --- | --- | --- |
| Flip Error | Beagle | 0.1977867 | 110.07309 | 0 | 0.1948284 | Inf |
| Flip Error | Eagle | 0.1595476 | 93.97320 | 0 | 0.1567524 | Inf |
| Flip Error | SHAPEIT | 0.1965362 | 115.14546 | 0 | 0.1937261 | Inf |
| Switch Error | Beagle | 0.0244927 | 32.94071 | 0 | 0.0232685 | Inf |
| Switch Error | Eagle | 0.0268967 | 36.95026 | 0 | 0.0256983 | Inf |
| Switch Error | SHAPEIT | 0.0227155 | 31.24358 | 0 | 0.0215185 | Inf |

**Table B2:** Average proportions of heterozygous positions and switch error locations by minor allele frequency bins in X chromosome synthetic diploids. Each synthetic diploid has a proportion of its heterozygous positions and its errors for each method within each of these bins, and here we present the average of these values over all 1000 synthetic diploids.

| Category | Rare | Uncommon | Common |
| --- | --- | --- | --- |
| Background | 0.0602 | 0.0853 | 0.855 |
| Beagle | 0.216 | 0.148 | 0.637 |
| Eagle | 0.317 | 0.142 | 0.541 |
| SHAPEIT | 0.202 | 0.144 | 0.654 |

**Table B3:** Average proportions of heterozygous positions and flip error locations by minor allele frequency bins in X chromosome synthetic diploids. Each synthetic diploid has a proportion of its heterozygous positions and its errors for each method within each of these bins, and here we present the average of these values over all 1000 synthetic diploids.

| Category | Rare | Uncommon | Common |
| --- | --- | --- | --- |
| Background | 0.0602 | 0.0853 | 0.855 |
| Beagle | 0.569 | 0.108 | 0.323 |
| Eagle | 0.56 | 0.112 | 0.328 |
| SHAPEIT | 0.562 | 0.112 | 0.326 |

**Table B4:**
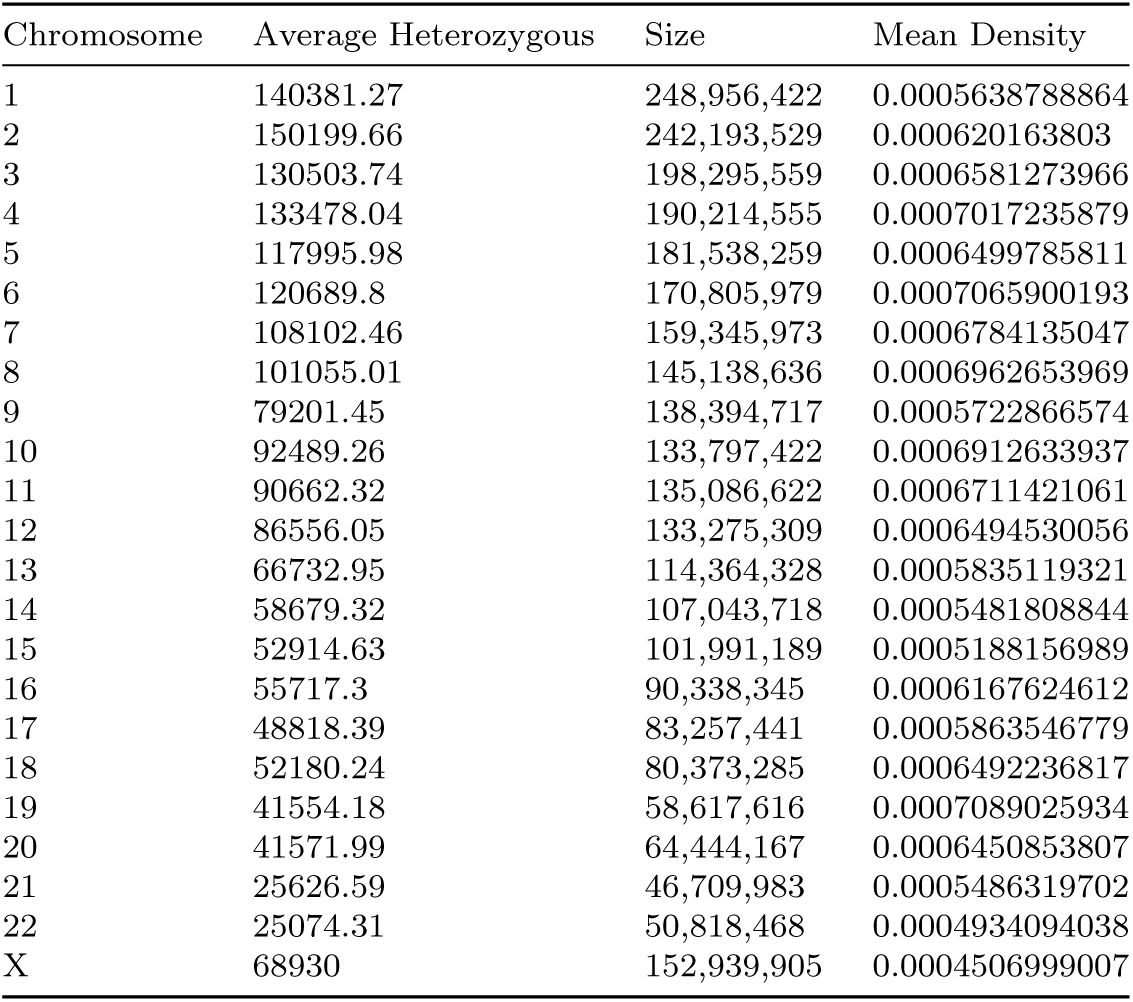
The mean number of heterozygous sites per proband (autosomes) and synthetic diploids (X) and chromosome sizes from GRCh38.

| Chromosome | Average Heterozygous | Size | Mean Density |
| --- | --- | --- | --- |
| 1 | 140381.27 | 248,956,422 | 0.0005638788864 |
| 2 | 150199.66 | 242,193,529 | 0.000620163803 |
| 3 | 130503.74 | 198,295,559 | 0.0006581273966 |
| 4 | 133478.04 | 190,214,555 | 0.0007017235879 |
| 5 | 117995.98 | 181,538,259 | 0.0006499785811 |
| 6 | 120689.8 | 170,805,979 | 0.0007065900193 |
| 7 | 108102.46 | 159,345,973 | 0.0006784135047 |
| 8 | 101055.01 | 145,138,636 | 0.0006962653969 |
| 9 | 79201.45 | 138,394,717 | 0.0005722866574 |
| 10 | 92489.26 | 133,797,422 | 0.0006912633937 |
| 11 | 90662.32 | 135,086,622 | 0.0006711421061 |
| 12 | 86556.05 | 133,275,309 | 0.0006494530056 |
| 13 | 66732.95 | 114,364,328 | 0.0005835119321 |
| 14 | 58679.32 | 107,043,718 | 0.0005481808844 |
| 15 | 52914.63 | 101,991,189 | 0.0005188156989 |
| 16 | 55717.3 | 90,338,345 | 0.0006167624612 |
| 17 | 48818.39 | 83,257,441 | 0.0005863546779 |
| 18 | 52180.24 | 80,373,285 | 0.0006492236817 |
| 19 | 41554.18 | 58,617,616 | 0.0007089025934 |
| 20 | 41571.99 | 64,444,167 | 0.0006450853807 |
| 21 | 25626.59 | 46,709,983 | 0.0005486319702 |
| 22 | 25074.31 | 50,818,468 | 0.0004934094038 |
| X | 68930 | 152,939,905 | 0.0004506999007 |

**Table B5:** Mean error rates observed in probands (autosomes) and synthetic diploids (X).

| Chromosome | Mean Error Rate |  |  |
| --- | --- | --- | --- |
|  | Beagle | Eagle | SHAPEIT |
| 1 | 0.0077 | 0.0075 | 0.0081 |
| 2 | 0.0077 | 0.0075 | 0.0082 |
| 3 | 0.0072 | 0.0070 | 0.0077 |
| 4 | 0.0067 | 0.0066 | 0.0072 |
| 5 | 0.0072 | 0.0070 | 0.0077 |
| 6 | 0.0065 | 0.0063 | 0.0069 |
| 7 | 0.0074 | 0.0073 | 0.0079 |
| 8 | 0.0073 | 0.0071 | 0.0078 |
| 9 | 0.0087 | 0.0085 | 0.0092 |
| 10 | 0.0073 | 0.0072 | 0.0077 |
| 11 | 0.0073 | 0.0071 | 0.0077 |
| 12 | 0.0074 | 0.0072 | 0.0079 |
| 13 | 0.0071 | 0.0070 | 0.0076 |
| 14 | 0.0078 | 0.0078 | 0.0084 |
| 15 | 0.0089 | 0.0088 | 0.0094 |
| 16 | 0.0090 | 0.0089 | 0.0095 |
| 17 | 0.0091 | 0.0091 | 0.0096 |
| 18 | 0.0076 | 0.0075 | 0.0081 |
| 19 | 0.0087 | 0.0086 | 0.0090 |
| 20 | 0.0089 | 0.0088 | 0.0093 |
| 21 | 0.0093 | 0.0092 | 0.0098 |
| 22 | 0.0105 | 0.0105 | 0.0109 |
| X | 0.0031 | 0.0037 | 0.0031 |

**Table B6:** Mean switch and flip error rates observed in probands (autosomes) and synthetic diploids (X).

| Chromosome | Mean Switch Rate |  |  | Mean Flip Rate |  |  |
| --- | --- | --- | --- | --- | --- | --- |
|  | Beagle | Eagle | SHAPEIT | Beagle | Eagle | SHAPEIT |
| 1 | 0.0019 | 0.0025 | 0.0019 | 0.0057 | 0.0050 | 0.0062 |
| 2 | 0.0018 | 0.0024 | 0.0018 | 0.0059 | 0.0051 | 0.0064 |
| 3 | 0.0017 | 0.0022 | 0.0017 | 0.0055 | 0.0048 | 0.0060 |
| 4 | 0.0015 | 0.0021 | 0.0015 | 0.0052 | 0.0045 | 0.0057 |
| 5 | 0.0017 | 0.0022 | 0.0016 | 0.0055 | 0.0048 | 0.0060 |
| 6 | 0.0015 | 0.0020 | 0.0014 | 0.0050 | 0.0044 | 0.0054 |
| 7 | 0.0019 | 0.0025 | 0.0019 | 0.0055 | 0.0048 | 0.0060 |
| 8 | 0.0017 | 0.0022 | 0.0017 | 0.0056 | 0.0049 | 0.0061 |
| 9 | 0.0026 | 0.0032 | 0.0025 | 0.0061 | 0.0053 | 0.0066 |
| 10 | 0.0019 | 0.0024 | 0.0018 | 0.0054 | 0.0048 | 0.0059 |
| 11 | 0.0017 | 0.0023 | 0.0017 | 0.0055 | 0.0048 | 0.0060 |
| 12 | 0.0018 | 0.0024 | 0.0018 | 0.0055 | 0.0048 | 0.0060 |
| 13 | 0.0018 | 0.0024 | 0.0018 | 0.0053 | 0.0046 | 0.0058 |
| 14 | 0.0021 | 0.0027 | 0.0021 | 0.0057 | 0.0050 | 0.0062 |
| 15 | 0.0027 | 0.0034 | 0.0027 | 0.0061 | 0.0054 | 0.0067 |
| 16 | 0.0026 | 0.0034 | 0.0026 | 0.0064 | 0.0055 | 0.0070 |
| 17 | 0.0028 | 0.0035 | 0.0027 | 0.0064 | 0.0056 | 0.0069 |
| 18 | 0.0021 | 0.0027 | 0.0021 | 0.0055 | 0.0048 | 0.0060 |
| 19 | 0.0027 | 0.0034 | 0.0026 | 0.0059 | 0.0052 | 0.0064 |
| 20 | 0.0028 | 0.0034 | 0.0027 | 0.0061 | 0.0054 | 0.0066 |
| 21 | 0.0032 | 0.0038 | 0.0032 | 0.0060 | 0.0054 | 0.0065 |
| 22 | 0.0038 | 0.0045 | 0.0037 | 0.0066 | 0.0060 | 0.0072 |
| X | 0.0017 | 0.0023 | 0.0016 | 0.0014 | 0.0014 | 0.0016 |

## Notes

### Competing Interest Statement

The authors have declared no competing interest.

### Summary of Updates

Some minor revisions to the text for style and clarity

